# A Feed-forward Pathway Drives LRRK2 kinase Membrane Recruitment and Activation

**DOI:** 10.1101/2022.04.25.489459

**Authors:** Edmundo G. Vides, Ayan Adhikari, Claire Y. Chiang, Pawel Lis, Elena Purlyte, Charles Limouse, Justin L. Shumate, Elena Spínola-Lasso, Herschel S. Dhekne, Dario R. Alessi, Suzanne R. Pfeffer

## Abstract

Activating mutations in the Leucine Rich Repeat Kinase 2 (LRRK2) cause Parkinson’s disease and previously we showed that activated LRRK2 phosphorylates a subset of Rab GTPases (Steger et al., 2017). Moreover, Golgi-associated Rab29 can recruit LRRK2 to the surface of the Golgi and activate it there for both auto- and Rab substrate phosphorylation. Here we define the precise Rab29 binding region of the LRRK2 Armadillo domain between residues 360-450 and show that this domain, termed “Site #1”, can also bind additional LRRK2 substrates, Rab8A and Rab10. Moreover, we identify a distinct, N-terminal, higher affinity interaction interface between LRRK2 phosphorylated Rab8 and Rab10 termed “Site #2”, that can retain LRRK2 on membranes in cells to catalyze multiple, subsequent phosphorylation events. Kinase inhibitor washout experiments demonstrate that rapid recovery of kinase activity in cells depends on the ability of LRRK2 to associate with phosphorylated Rab proteins, and phosphorylated Rab8A stimulates LRRK2 phosphorylation of Rab10 in vitro. Reconstitution of purified LRRK2 recruitment onto planar lipid bilayers decorated with Rab10 protein demonstrates cooperative association of only active LRRK2 with phospho-Rab10-containing membrane surfaces. These experiments reveal a feed-forward pathway that provides spatial control and membrane activation of LRRK2 kinase activity.

## Introduction

Activating mutations in the Leucine Rich Repeat Kinase 2 (LRRK2) cause inherited Parkinson’s disease, and lead to the phosphorylation of a subset of Rab GTPases (Alessi and Sammler, 2018), in particular, Rab8A, Rab10, and Rab29 within a conserved residue of the Switch-II effector binding motif. Rab GTPases are master regulators of membrane trafficking and are thought to serve as identity determinants of membrane bound compartments of the secretory and endocytic pathways (Pfeffer, 2017). In their GTP bound forms, Rabs are best known for their roles in linking motor proteins to transport vesicles and facilitating the process of transport vesicle docking.

Our previous work showed that Rab phosphorylation blocks the ability of Rab proteins to be activated by their cognate guanine nucleotide exchange factors or to bind to the GDI proteins that recycle GDP-bearing Rabs from target membranes to their membranes of origin (Steger et al., 2016; 2017). Moreover, phosphorylation of Rab8A and Rab10 blocks their ability to bind known effector proteins and enhances binding to a novel set of effectors that includes RILPL1, RILPL2, JIP3, JIP4 and MyoVa proteins (Steger et al., 2017; Waschbüsch et al., 2020; Dhekne et al., 2021). Thus, Rab phosphorylation flips a switch on Rab effector selectivity that can drive dominant physiological changes, including blocking primary cilia formation (Steger et al., 2017; Dhekne et al., 2018; Sobu et al., 2021; Khan et al., 2021) and autophagosome motility in axons (Boecker et al., 2021).

Most LRRK2 is found in the cell cytosol where it appears to be inactive (Biskup et al., 2006; Berger et al., 2010; Purlyte et al., 2018). Recent structural analysis of the catalytic, C-terminal half of LRRK2 (Deniston et al., 2021) and full length human LRRK2 protein yielded structures of both monomeric and dimeric, inactive states (Myasnikov et al., 2021). Several groups have reported that active LRRK2 is a dimer (Greggio et al., 2008; Klein et al., 2009; Sen et al., 2009; Berger et al, 2010; Civiero et al., 2012; Guaitoli et al, 2016), and higher order forms were detected on membranes upon crosslinking (Berger et al., 2010; Schapansky et al., 2014) and upon Rab29 binding (Zhu et al., 2022). Thus, LRRK2 membrane association is associated with kinase activation, however the molecular basis for this activation is not yet known.

Exogenously expressed, Golgi-localized Rab29 protein can recruit LRRK2 onto membranes and activate it there for both auto- and Rab substrate phosphorylation (Kuwahara et al., 2016; Liu et al., 2018; Purlyte et al., 2018; Madero-Pérez et al., 2018). Indeed, even Rab29 artificially anchored on mitochondria can activate LRRK2 and drive its membrane recruitment (Gomez et al., 2019). McGrath et al. (2021) implicated LRRK2 residues 386-392 as being important for the interaction of Rab29/32/38 family members with the LRRK2 kinase Armadillo domain. However, the LRRK2 Armadillo domain is located at some distance from the kinase domain, at least in the current structure models for LRRK2 protein (Myasnikov et al., 2021). Thus, how Rab29 binding might activate LRRK2 kinase activity is not at all clear. In addition, because Rab29 is not needed for LRRK2 action on Rab8A or Rab10 proteins (Kalogeropulou et al., 2020), other pathways for LRRK2 activation must exist.

In this study, we define a specific patch (“Site #1”) of the LRRK2 Armadillo domain that binds to Rab8A, Rab10 and Rab29 protein with affinities similar to those reported previously (McGrath et al., 2021). More importantly, we identify a distinct region of LRRK2 Armadillo domain (“Site #2”) that binds specifically to LRRK2-*phosphorylated* Rab8A and Rab10 proteins, to establish a feed forward activation mechanism for membrane-associated LRRK2 kinase.

## Results

### Rab29 binds to the C-terminal portion of the LRRK2 Armadillo domain

McGrath et al. (2021) showed that the LRRK2 Armadillo domain residues 1-552 contain a binding site that interacts specifically with purified Rab29, 32 and 38 in vitro with affinities of 2.7, 1.2 and 1.2-2.4µM, respectively. We used microscale thermophoresis to determine the affinity of other Rab GTPase substrates with this portion of LRRK2 kinase. For these experiments, portions of the LRRK2 Armadillo domain were fluorescently labeled and incubated with Rab GTPases in the presence of Mg^2+^-GTP. Figure 1 shows binding curves for Rab29 with full length Armadillo domain (residues 1-552, panel A), as well as sub-fragments composed of LRRK2 residues 1-159 (Fig. 1C) or 350-550 (Fig. 1D). Rab29 showed specific binding to the full length 1-552 Armadillo fragment with a K_D_ of 1.6µM (Fig. 1A), comparable to that reported previously using other methods (McGrath et al., 2021). Under these conditions, the non-LRRK2 substrate Rab7 protein failed to bind to the Armadillo 1-552 fragment (Fig. 1B). No Rab29 binding was detected to a fragment representing the N-terminal 1-159 LRRK2 residues (binding >29µM; Fig. 1C); essentially full binding was observed with a fragment encompassing residues 350-550 (K_D_ = 1.6µM; Fig. 1D). Thus, Rab29 binds to the C-terminal portion of LRRK2’s Armadillo domain at a site that we will refer to as Site #1.

**Figure 1.**
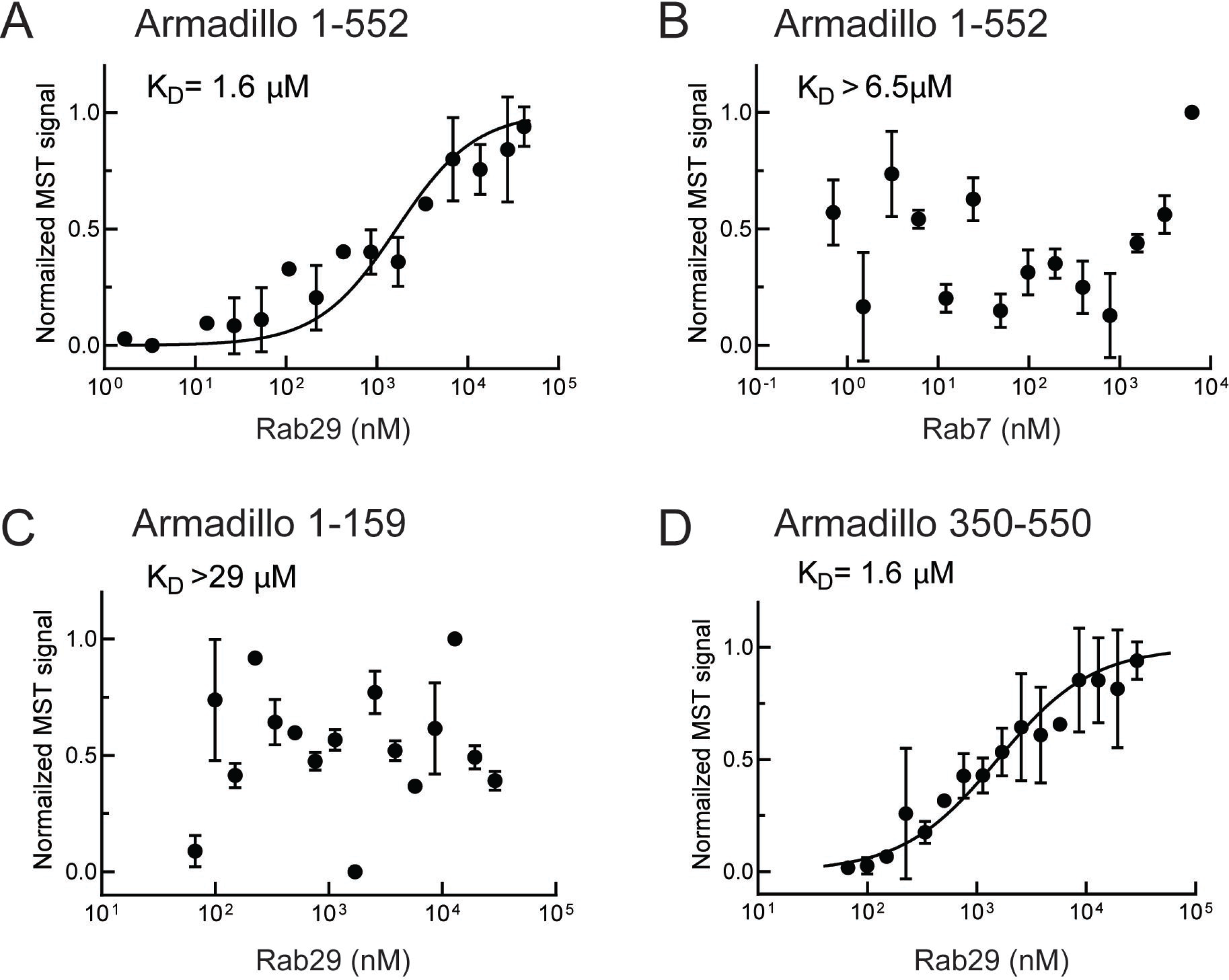
Rab29 binds to the C-terminal portion of the LRRK2 Armadillo domain. Microscale thermophoresis of full length (residues 1-552), labeled LRRK2 Armadillo domain with His-Rab29 (A) or with His-Rab7 (B). (C, D) Microscale thermophoresis of labeled LRRK2 Armadillo domain residues 1-159 (C) or 350-550 (D) with Rab29. Purified Rab29 was serially diluted and then NHS-RED labeled-LRRK2 Armadillo (final concentration 100nM) was added. Graphs show mean and SEM from three independent measurements, each from a different set of protein preparations.

### Rab8A and Rab10 bind to the LRRK2 Armadillo domain

Similar experiments were carried out with Rab8A and Rab10, the most prominent LRRK2 substrates (Steger et al., 2017). Rab8A bound full length Armadillo domain with a K_D_ of 2.9µM (Fig. 2A), showed weaker interaction with the LRRK2 1-159 fragment (K_D_ ∼ 6.7µM; Fig. 2B), and good binding to the 350-550 fragment (K_D_ = 2.3µM; Fig. 2C). These data indicate that Rab8A may bind to the same site as Rab29. Like Rab8A, Rab10 bound to full length Armadillo 1-552 with a K_D_ of 2.4µM (Fig. 2D); weaker binding was detected for 1-159 and 350-550 fragments, yielding K_D_s of 5.1µM in both cases (Fig. 2E,F). Thus, in addition to Rab32, 38 and 29, Rabs 8A and 10 can bind to LRRK2 residues 350-550. Note that Rab32 and Rab38 are not substrates of LRRK2 kinase as they lack a phosphorylatable Ser/Thr residue in the Switch-II motif (Steger et al., 2016; 2107); they show extremely narrow tissue-specific expression but are related to Rab29 protein.

**Figure 2.**
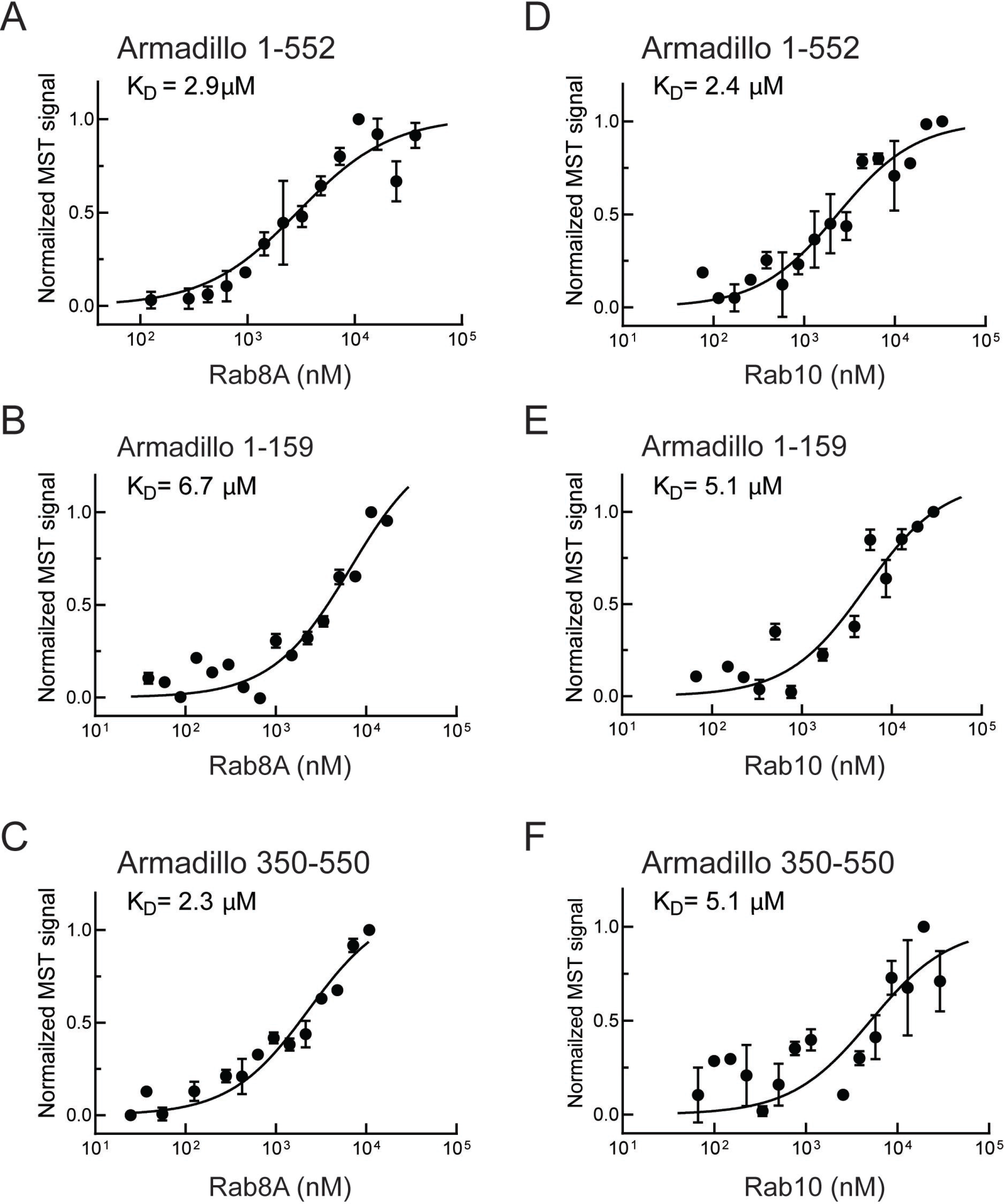
Rab8A and Rab10 bind to the LRRK2 Armadillo domain. (A-C) Microscale thermophoresis of labeled, LRRK2 Armadillo domain fragments comprised of residues 1-552, 1-159, or 350-550 with Rab8A Q67L (1-181) as indicated. (C, D, E) Microscale thermophoresis for Rab10 Q68L (1-181) with indicated LRRK2 Armadillo fragments, as in A. Purified Rab proteins were serially diluted and then NHS-RED labeled LRRK2 Armadillo domain (final concentration 100nM) was added. Graphs show mean and SEM from three independent measurements, each from a different set of protein preparations.

### Residues critical for Rab GTPase binding to LRRK2 residues 350-550, Site #1

Previous work implicated LRRK2 residues 386–392 in contributing to a Rab29/32/38 binding interface (McGrath et al., 2021). We used a microscopy-based assay to identify any portions of the first 1000 residues of LRRK2 that would relocalize to the Golgi upon co-expression with Golgi localized, HA-Rab29 protein (Fig. 3-Fig. Suppl. 1). Twenty two constructs were transfected into cells and their localization scored visually. The smallest fragment of LRRK2 that interacted with HA-tagged Rab29 in HeLa cells, thereby co-localizing at the Golgi complex, encompassed LRRK2 residues 350-550.

We next deployed AlphaFold docking (Jumper et al., 2021) using ColabFold (Mirdita et al., 2022) and the AlphaFold2_advanced.ipynb notebook with the default settings to model the interaction of Rab29 with the LRRK2 350-550 fragment (Fig. 3A and Fig. 3-Fig. Suppl. 2). Residues highlighted in red show key contacts between LRRK2 and Rab29 and will be shown below to be essential for detection of this interaction in cells. This modelled structure of Site #1 is extremely similar to that of the recently reported experimental cryo-EM structure of Rab29 complexed full length LRRK2 (Zhu et al 2022).

**Figure 3.**
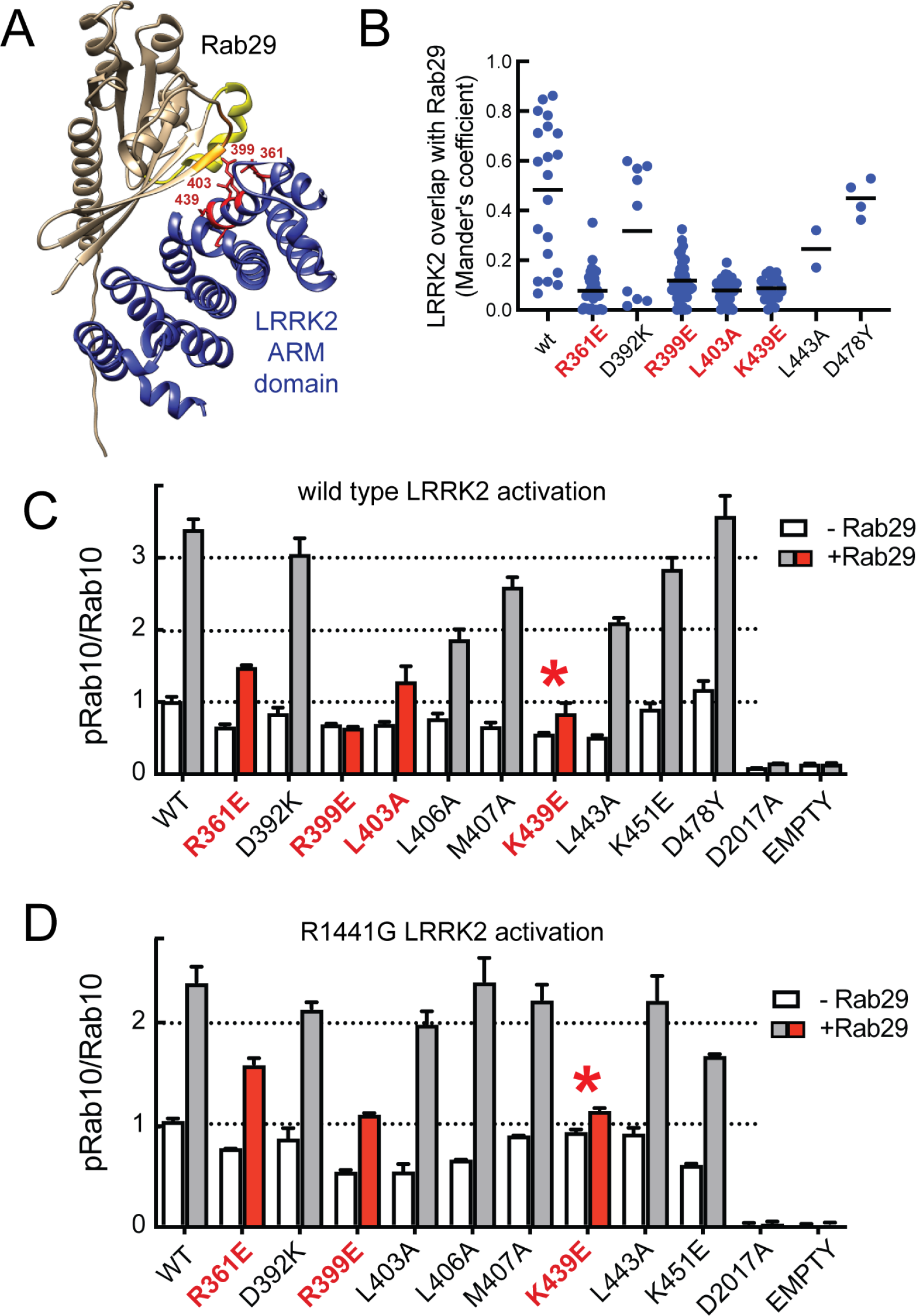
Characterization of critical LRRK2 residues mediating binding to Rab29. (A) Predicted interactions between Rab29 and the LRRK2 Armadillo domain using AlphaFold docking (Jumper et al., 2021), ColabFold (Mirdita et al., 2021) and the AlphaFold2_advanced.ipynb notebook default settings. Residues identified in red show key contacts between LRRK2 and Rab29; orange and yellow coloring indicates the Switch I and Switch II domains of Rab29. (B) The wild type and indicated mutants of full length of GFP-LRRK2 were co-expressed with HA-Rab29 in HeLa cells. 24h post transfection cells were fixed and localization assessed by confocal microscopy. LRRK2 overlap with Rab29 is presented as a Mander’s coefficient determined using Cellprofiler software (McQuin et al., 2018). (C, D) Wild type and indicated mutants of full length of GFP-LRRK2 (C) or GFP-LRRK2 R1441G (D) were co-expressed with HA-Rab29 in HEK293T cells. 24h post transfection, cells were lysed and extracts immunoblotted with the indicated antibodies. Shown are the averages and standard deviations of duplicate determinations; red asterisks indicate preferred mutant.

Three metrics were used to evaluate the importance of individual residues to contribute to Rab29 interaction. First, we tested the impact of mutations on the ability of full length LRRK2 to co-localize with HA-Rab29 at the Golgi in HeLa cells (Fig. 3B and Fig.3 – Fig. Suppl. 3); we also tested the ability of exogenously expressed Rab29 to stimulate activity of the same point mutants in the background of either wild type LRRK2 (Fig. 3C, Fig. 3–Fig. Suppl. 4A) or pathogenic R1441G LRRK2 (Fig. 3D, Fig 3.--Fig. Suppl. 4B). This work identified 4 key mutations of highly conserved residues (R361E, R399E, L403A and K439E) that blocked both the co-localization of LRRK2 and Rab29 in HeLa cells (Fig. 3B, red) as well as activation of LRRK2 upon overexpression of Rab29 in HEK293 cells (Fig. 3C, red). In experiments undertaken with pathogenic R1441G LRRK2 that is more potently activated by Rab29, the K439E LRRK2 mutation completely blocked LRRK2 kinase activation; R399E showed weak activation (Fig. 3D and Fig. 3--Fig. Suppl. 4B). Some of the other mutants blocked co-localization with Rab29 in HeLa cells without completely suppressing LRRK2 activation following overexpression of Rab29. We therefore recommend using the Site#1 K439E LRRK2 mutation to block Rab29 interaction and activation in future work (asterisks in Figs. 3B,C) as it shows the lowest amount of Rab29 activation with pathogenic R1441G LRRK2. Altogether, these data highlight the importance of a surface that is comprised of LRRK2 residues Arg361, Arg399, Leu403, Lys439 in binding Rab GTPases (Site #1) (Fig. 3A and Fig. 3 -Fig. Suppl. 2). Analysis of Rab8A interaction with the LRRK2 350-550 fragment using AlphaFold within Chimera X 1.4 confirmed the importance of the same LRRK2 residues for Rab8A interaction in silico (Figure 3-Fig. Supplement 2C).

### PhosphoRab binding to LRRK2, Site #2

To understand the consequences of LRRK2-mediated Rab GTPase phosphorylation, it is important to identify specific binding partners of phosphorylated Rab proteins, and to study the consequences of such binding events. We recently established a facile method that enables us to monitor phosphoRab binding to proteins of interest in conjunction with microscale thermophoresis binding assays. Briefly, Rab proteins are phosphorylated >90% in vitro by MST3 kinase (Dhekne et al., 2021; Vides and Pfeffer, 2021) that phosphorylates Rab proteins at the same position as LRRK2 kinase (Vieweg et al., 2020). Recombinant MST3 is much easier to purify in large amounts for biochemical experiments than LRRK2. We used this assay to monitor the possible interaction of phosphorylated LRRK2 substrates to the LRRK2 Armadillo domain and were delighted to discover that pRab8A and pRab10 proteins bind with high affinity to a site distinct from that used by non-phosphorylated Rab proteins that we term site #2.

As shown in Fig. 4, phosphoRab8A and phosphoRab10 bound with K_D_s of ∼900nM and 1µM to the full Armadillo domain 1-552 fragment, respectively (Figs. 4A, D); this binding reflected interaction with N-terminal LRRK2 residues 1-159, as this fragment was sufficient to yield essentially the same K_D_s of 1µM and 700 nM, respectively for phosphoRab8A and phosphoRab10 proteins (Figs. 4B, E). Furthermore, no binding was detected for phosphoRab8A or phosphoRab10 with LRRK2 residues 350-550 (Figs. 4C, F). These data demonstrate that Rab8A and Rab10 GTPases, phosphorylated at the same residues modified by LRRK2 kinase bind very tightly to the LRRK2 N-terminus but no longer interact with the 350-550 region that interacts with dephosphorylated Rab proteins.

**Figure 4.**
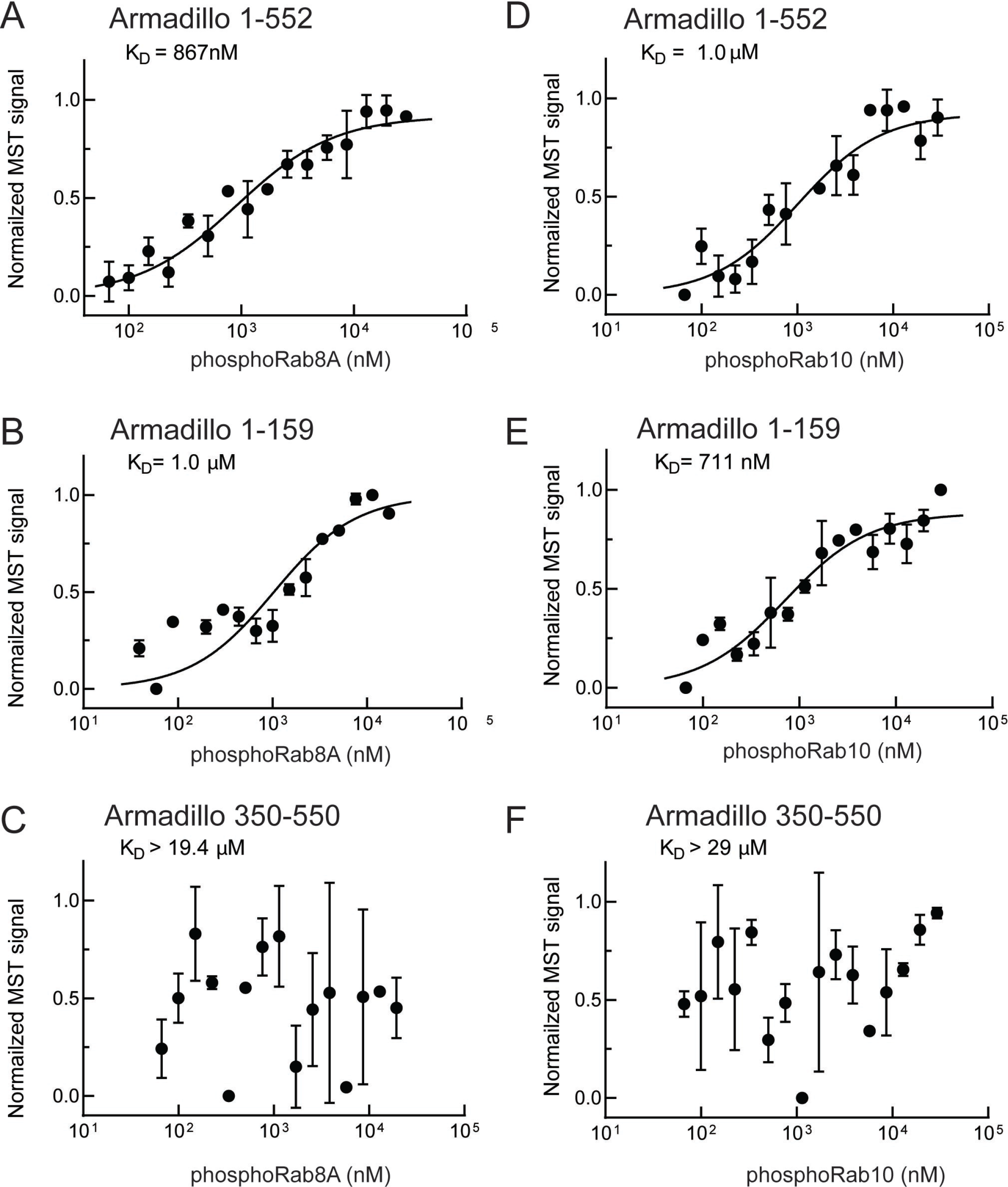
phosphoRab8A and phosphoRab10 bind with high affinity to the N-terminal portion of the LRRK2 Armadillo domain. (A-F) Microscale thermophoresis of labeled, indicated, LRRK2 Armadillo fragments with His-phosphoRab8A Q67L 1-181 (A-C) or with His phosphoRab10 Q68L 1-181 (pRab10 (D-F). Purified Rab proteins were phosphorylated with Mst3 kinase at 27°C for 2 h and then serially diluted; NHS-RED labeled Armadillo (final concentration 100 nM) was then added. Graphs show mean and SEM from three independent measurements, each from a different set of protein preparations.

Note that non-phosphorylated Rab8A and Rab10 also bound to the Site #2-containing fragment 1-159 with relatively weak affinities of 5 or 6µM (Fig. 2B,E; Table 1). Interestingly, AlphaFold in Chimera X predicts that the 1-159 fragment contains a potential, non-phosphoRab binding site that is occluded in a longer fragment (1-400), and thus also in full length LRRK2. Moreover, as discussed below, these K_D_ values may be higher than the concentrations of these Rab GTPases in cells, thus it seems unlikely that non-phosphoRabs interact with Site #2 under normal physiological conditions. We conclude that phosphoRab binding is the predominant interaction between LRRK2 1-159 and Rab GTPases.

**Table 1.** Summary of Binding Affinities. Note that these values are likely underestimates of affinities as typical preparations of purified Rab proteins contained ∼50% bound GDP and ∼50% bound GTP by mass spectrometry. Non-phosphorylated Rab interaction with Armadillo 1-159 is shown in parentheses as it likely reflects binding to an Alphafold-predicted site near the C-terminus of this fragment that will not be accessible in full length LRRK2 protein.

|  | Armadillo<br>1-159<br>(Site #2-containing) | Armadillo<br>1-552 | Armadillo<br>350-550<br>(Site #1-containing) | Armadillo<br>1-552<br>K17A | Armadillo<br>1-552<br>K18A |
| --- | --- | --- | --- | --- | --- |
| Rab29 | >29 | 1.6 ± 0.9 | 1.6 ± 0.5 | - | - |
| Rab10-Q68L | (5.1 ± 3.1) | 2.4 ± 0.6 | 5.1 ± 2.5 | - | - |
| pRab10-Q68L | 0.71 ± 0.3 | 1.0 ± 0.4 | >29 | >20 | >20 |
| Rab8A-Q67L | (6.7 ± 3.6) | 2.9 ± 1.2 | 2.3 ± 1.0 | - | - |
| pRab8A-Q67L | 1.0 ± 0.6 | 0.87 ± 0.4 | >19.4 | - | - |
| Rab7 | - | >6.5 | - | - | - |

Electrostatic analysis of a model of the LRRK2 Armadillo domain revealed that the absolute N-terminus of LRRK2 contains a patch of basic amino acids (highlighted in blue) that may comprise a phosphoRab interaction interface (Fig. 5A) (Jurus et al., 2018; Pettersen et al., 2004). Such modeling led us to test the role of lysine residues at positions 17 and 18 in mediating LRRK2 interaction. Mutation of either lysine 17 or 18 abolished phosphoRab10 binding to LRRK2 Armadillo domain, with binding decreased to >20µM upon single mutation at either site (Fig. 5C, D). When the conservation score of these residues is analyzed using the Consurf server (Ashkenazy et al., 2016) K17 and K18 have a score or 2 and 8 respectively (9 is the maximum score), indicating that K18 is highly conserved and plays an especially important role. These experiments define a second, Rab binding site #2 that is specific for phosphorylated Rab proteins (Fig. 5B).

**Figure 5.**
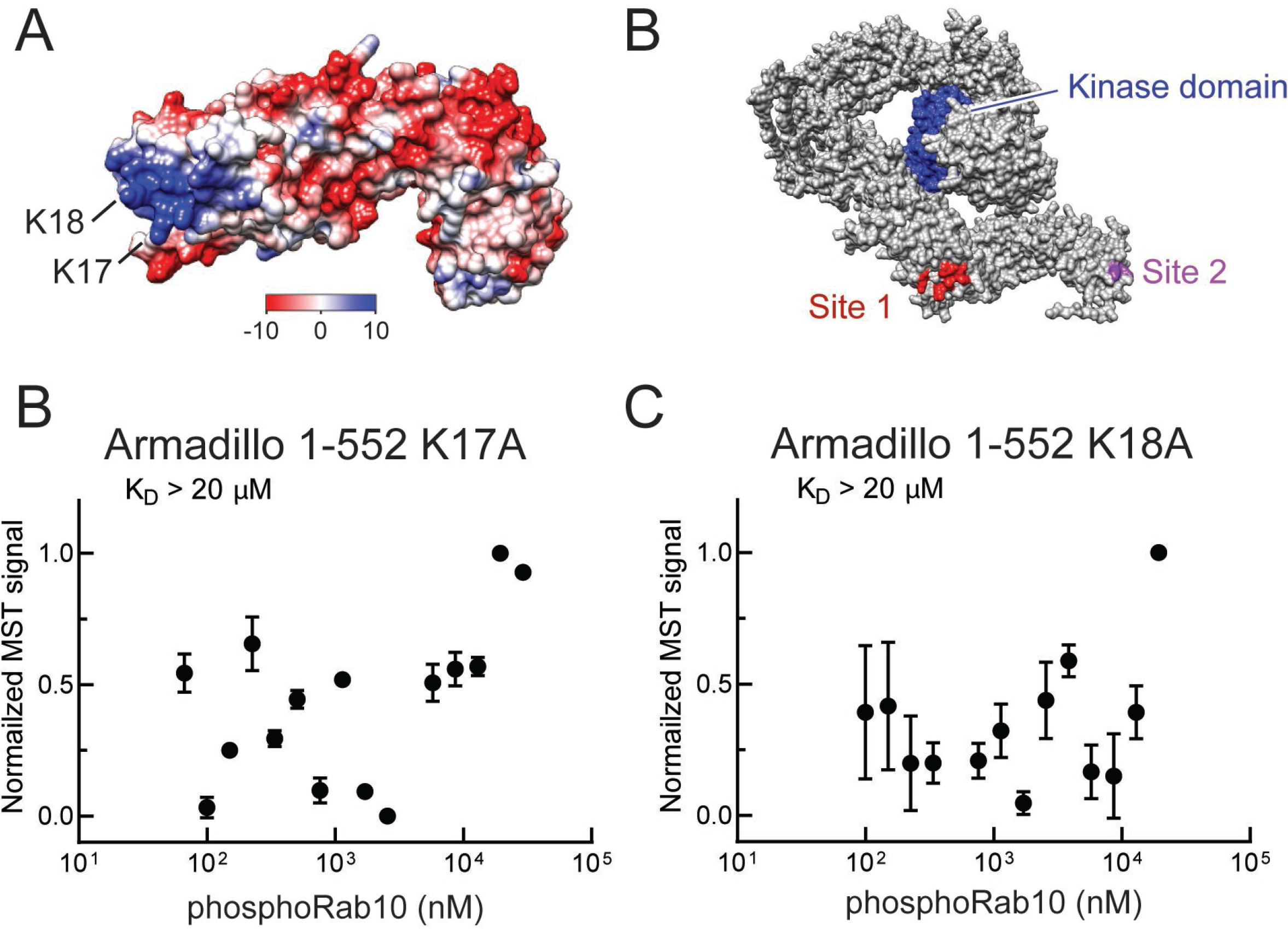
A. Electrostatic surface potential of LRRK2 Armadillo domain residues 1-552 modeled using Chimera 2 software (Petterson et al., 2004); blue indicates a positively charged surface. LRRK2 K17 and K18 are indicated. (B) Alphafold (Jumper et al., 2021) structure of putative, active LRRK2 with residues that mediate Rab29 binding shown in red (Site #1) and the K17/K18 residues that are required for phosphoRab10 binding (Site #2) shown in magenta; the kinase domain is shown in blue. (C,D) Microscale thermophoresis of labeled, full length LRRK2 K17A or K18A Armadillo 1-552 with His phosphoRab10 Q68L 1-181. Purified Rab10 protein was phosphorylated with Mst3 kinase at 27°C for 2 h and then serially diluted; NHS-RED labeled Armadillo (final concentration 100 nM) was then added. Graphs show mean and SEM from three independent measurements, each from a different set of protein preparations.

To determine the significance of the phosphoRab binding site in relation to LRRK2 membrane recruitment in cells, we generated full length FLAG-LRRK2 protein containing point mutations at both lysines 17 and 18 and investigated its cellular localization upon expression in HeLa cells (Fig. 6). To improve our ability to detect membrane associated LRRK2 distribution, cells grown on collagen coated coverslips were dipped in liquid nitrogen and then thawed in a physiological, glutamate-containing buffer to crack open the plasma membrane and release cytosolic proteins prior to fixation (Seaman et al., 2004; Purlyte et al., 2018). Under these conditions, LRRK2 co-localizes with phosphorylated Rab proteins (Purlyte et al., 2018; Sobu et al., 2021).

**Figure 6.**
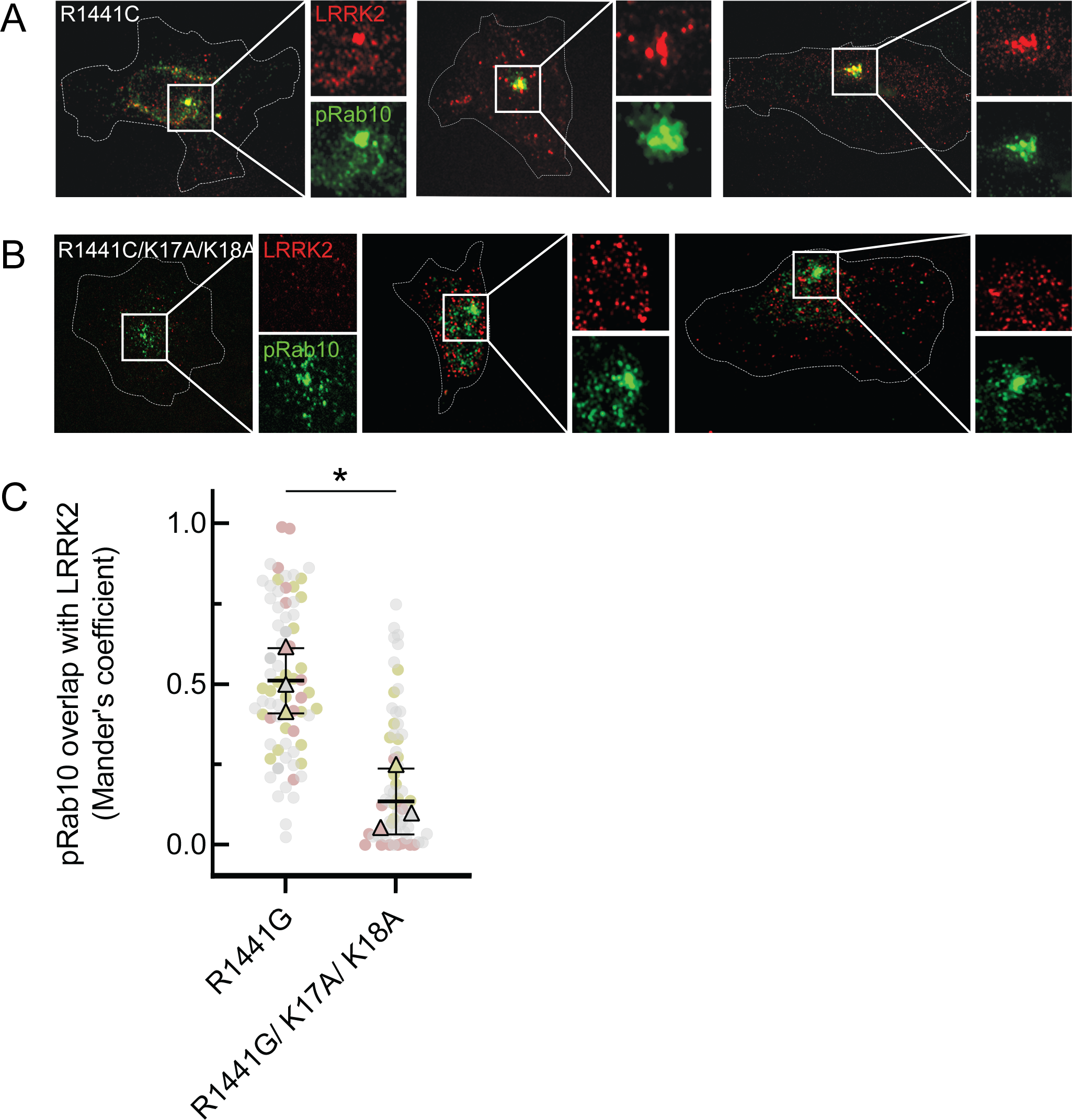
LRRK2 K17 and K18 are critical for pRab10 interactions in cells. (A) FLAG-LRRK2 R1441G (red) was transfected into HeLa cells plated on collagen coated coverslips and co-localized with endogenous wild type pRab10 (green). Cells on coverslips were dipped in liquid nitrogen to deplete cytosol and enhance membrane-bound signal. Insets show enlargements of boxed areas representing peri-centriolar LRRK2 and pRab10. (B) FLAG-LRRK2 R1441G/K17A/K18A (red) was transfected into HeLa cells plated on collagen coated coverslips and stained and localized with pRab10 (green) as in A. (C) Quantification of pRab10 overlap with LRRK2 by Mander’s coefficient. Error bars represent SEM of means from three different experiments (represented by colored dots), each with >40 cells per condition. Significance was determined by *t* test, *P = 0.0108.

As expected, PhosphoRab10 was detected as a bright spot adjacent to the mother centriole in HeLa cells (Fig. 6A), and the co-expressed, R1441G pathogenic mutant LRRK2 protein showed good co-localization with phosphoRab10 protein (Figs. 6A,C), as we have reported previously (Purlyte et al., 2018; Sobu et al., 2021). In contrast, although exogenously expressed, R1441G LRRK2 bearing K17/18/A mutations still generated a perinuclear, phosphoRab10-containing structure, LRRK2 displayed much less co-localization with the phosphoRab proteins or with membranes overall (Figs. 6B,C). These experiments show that K17 and K18 are important for exogenous LRRK2 membrane association with a pool of highly phosphorylated Rab10 protein. The importance of LRRK2’s N-terminal lysine residues also suggests that caution may be in order when evaluating membrane interactions of LRRK2 tagged N-terminally with larger tags such as GFP that may hinder access to K17/K18.

### PhosphoRab-LRRK2 interaction increases rates of kinase recovery

We next explored the relevance of phosphoRab binding to LRRK2’s N-terminus in relation to the overall kinetics of Rab phosphorylation in cells. LRRK2-mediated Rab GTPase phosphorylation is a highly dynamic process that is counteracted by the action of PPM1H phosphatase (Berndsen et al., 2019); at steady state, only a small fraction of total Rab proteins are LRRK2-phosphorylated (Ito et al., 2016). The initial rate of kinase activity can be determined by monitoring the phosphorylation of Rab10 protein after washout of the LRRK2 inhibitor, MLi-2 (Ito et al., 2016; Kalogeropulou, 2020).

When HeLa cells were treated with 200nM MLi-2 for 1hr and then washed with culture medium, Rab10 was efficiently re-phosphorylated by exogenous, FLAG-tagged, R1441G LRRK2 protein over the 2h time course evaluated (Fig. 7A, E). In contrast, cells expressing FLAG-R1441G LRRK2 bearing K17/18A mutations showed comparable total phosphoRab10 levels to begin with, but significantly slower re-phosphorylation (Fig 7B, E). Similar results were obtained in experiments comparing the re-activation of FLAG-tagged, wild type LRRK2 (Fig. 7C,F) with that of LRRK2 K17/18A (Fig. 7D,F). As reported previously (Ito et al., 2016), wild type LRRK2 recovery was more efficient than that of R1441G LRRK2. In summary, these experiments demonstrate that K17/K18 residues are important for efficient reactivation of LRRK2 after MLI-2 washout, consistent with their role in anchoring LRRK2 at sites adjacent to phosphorylation substrates.

**Figure 7.**
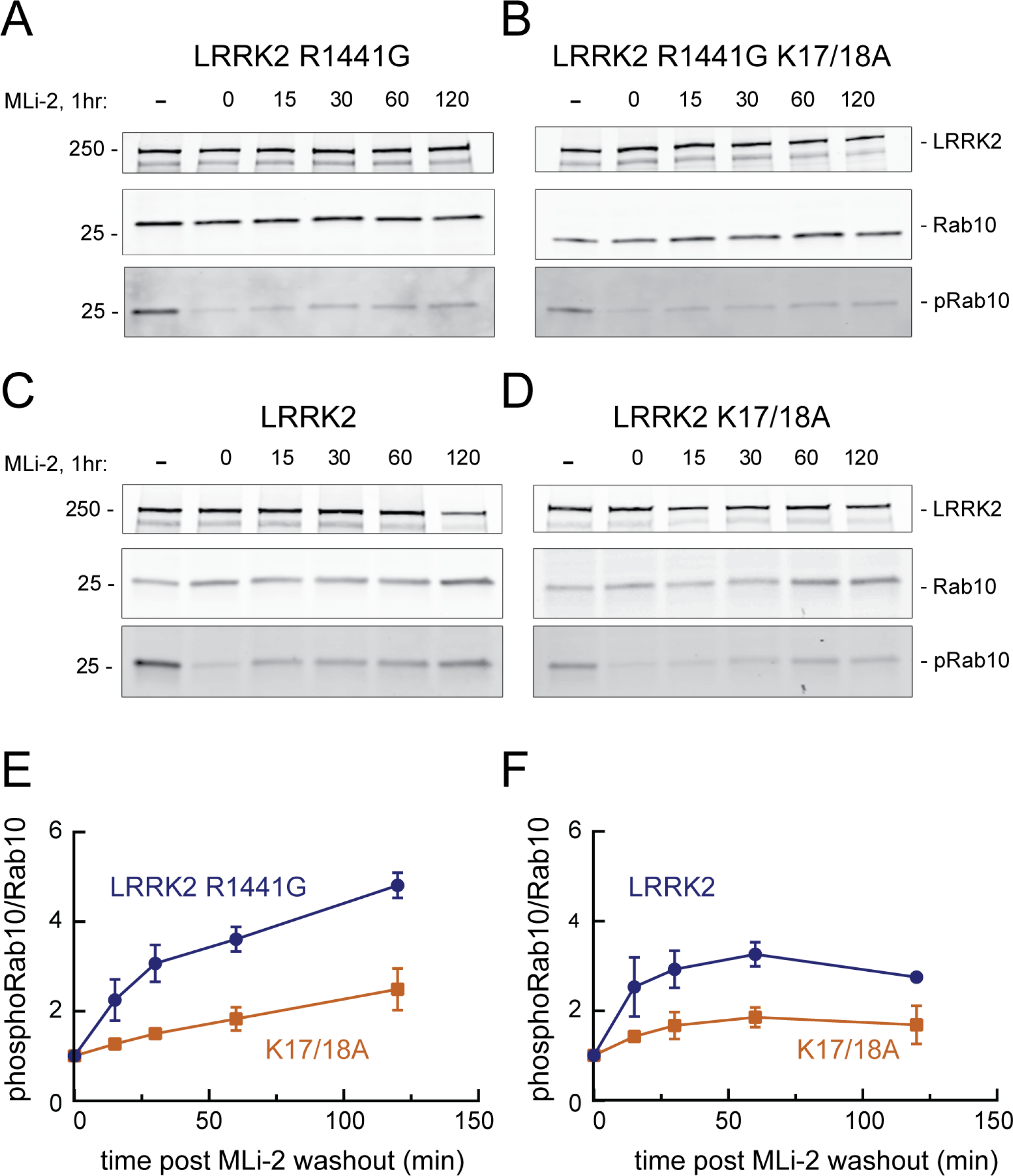
LRRK2 K17 and K18 increase endogenous pRab10 recovery after LRRK2 inhibitor washout. (A-D) FLAG-LRRK2 R1441G, FLAG-LRRK2 R1441G/K17A/K18A, LRRK2, or LRRK2 K17A/K18A was transfected into HeLa cells. 48 hr post transfection cells were treated with 200nM of MLi-2 for 1 hr. The MLi-2 was then removed by multiple washes and incubated for the indicated times prior to cell lysis. Whole cell extracts (20µg) were subjected to quantitative immunoblot analysis using anti-LRRK2, anti-Rab10, and anti-pRab10 antibodies. (E-F) Quantification of pRab10/ total Rab10 fold change and normalized to no MLi2 control. Error bars represent mean ± SD from 2 different experiments per condition.

### Cooperative LRRK2 membrane recruitment on Rab-decorated planar lipid bilayers

Binding of phosphoRabs to Site #2 at the N-terminus of LRRK2 (Fig. 5B) would set up a feed-forward process whereby the product of an initial phosphorylation reaction would enhance subsequent Rab GTPase phosphorylation by holding the enzyme on the surface of membranes that contain relevant Rab GTPase substrates. To visualize the membrane association process directly, we established a planar lipid bilayer system that would enable us to monitor the interaction of fluorescently labeled, purified, full length LRRK2 kinase with membrane anchored Rab10 substrate (Adhikari et al., 2022). For this purpose, bilayers were formed on the surface of glass bottom chambers comprised of phospholipids of a composition similar to that found in the Golgi (65% DOPC, 29% DOPS, 1% PI(4)P (Thomas and Fromme, 2016), mixed with 0.1% of the lipophilic tracer DiD dye and 5% DOGS-NTA [Ni^2^] to to enable anchoring of C-terminally His-GFP-tagged Rab10 protein. Binding of fluorescently labeled, hyperactive R1441G LRRK2 was then visualized in real time using total internal reflection (TIRF) light microscopy. Reactions were carried out in the presence of ATP, GTP and an ATP regenerating system to provide physiological conditions for the full length LRRK2 enzyme. Note that we routinely utilize R1441G LRRK2 because it is a highly active kinase in cells, although in vitro, R1441G LRRK2 displays the same level of Rab kinase activity as wild type LRRK2 (cf. Steger et al., 2017).

As shown in Figure 8A (red dots), fluorescent R1441G LRRK2 bound efficiently to lipid bilayers, only in the presence of pre-anchored Rab10 protein (compare with purple dots in 8B) and not when Rab11 protein was instead employed (Fig. 8B, green dots; movies 1-3). Importantly, almost no binding was observed with kinase inactive D2017A LRRK2 (Fig. 8A yellow dots, movie 4) (Steger et al., 2016). This indicates that at least Rab10 GTPase binding to Site #1 residues 361-451 results in a low affinity interaction that is not sufficient to retain this inactive LRRK2 protein on the bilayer under these conditions (7nM LRRK2, 2.5 µM Rab10). Reactions containing the Type I MLi-2 inhibitor showed aggregation of the fluorescent LRRK2 protein, as has been seen in cells. Incubations containing the Type 2 inhibitor, GZD-824 (Tasegian et al., 2021) showed weak binding, consistent with a requirement for phosphoRab10 generation to support LRRK2 binding to Site #2’s K17 and K18; however, under these conditions, LRRK2 was not monodisperse and could not be analyzed further. Importantly, R1441G LRRK2 mutated at lysines 17 and 18 bound to a lower extent than R1441G LRRK2 (Fig. 8A, blue dots; movie 5), confirming their important role in binding to phosphorylated Rab8A and Rab10. It is noteworthy that the K17/K18 mutant protein showed higher binding than the D2017A mutant, suggesting that a non-phosphoRab binding site may be more accessible for binding in an active versus inactive LRRK2 protein conformation.

**Figure 8.**
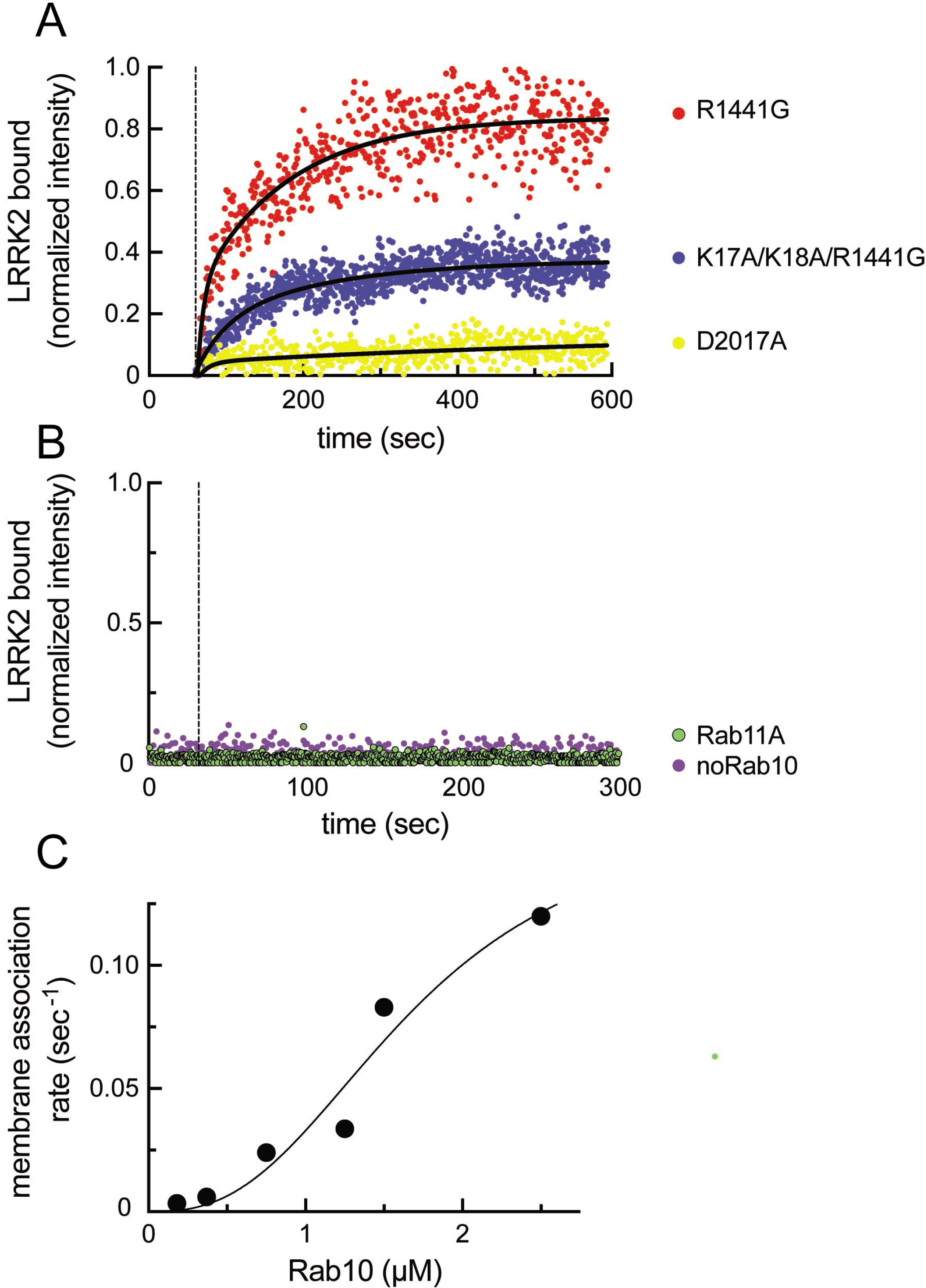
Feed-forward pathway for Rab10 phosphorylation is dependent on LRRK2 kinase activity. (A) Fluorescence intensity traces of individual, single molecules of 7nM CF633-labeled FLAG-LRRK2 R1441G on a substrate supported lipid bilayer decorated with lipid anchored GFP-Rab10 Q68L-His across 600 sec of live TIRF microscopy. Red, R1441G; Blue, K17A/K18A/R1441G; Yellow, D2017A. B. Reactions were carried out as in (A) except Rab10 was omitted (purple) or Rab10 was replaced with Rab11 (green). Dashed lines in A and B represent time of addition of fluorescently labeled LRRK2 at 60 sec. Fluorescence intensity was fitted by a nonlinear regression curve for two phase association. Fold change was calculated by dividing the average fluorescence intensity at steady state and subtracting background fluorescence intensity average determined from 60 sec prior to LRRK2 addition. C. Rate of membrane association of LRRK2 as a function of Rab10 concentration. This curve was fitted by a nonlinear regression fit using PRISM software (Mathworks) to determine a Hill coefficient.

Analysis of the kinetics of LRRK2 binding as a function of Rab protein concentration showed clear, cooperative membrane association of R1441G LRRK2, consistent with a feed-forward mechanism, as predicted from the in vitro Rab binding data (Fig. 8C). A nonlinear regression fit of the data indicated a Hill coefficient of 2.7, consistent with a positive, cooperative phenomenon. In summary, these data demonstrate that LRRK2 kinase is recruited to membranes and then held there by phosphorylated Rabs to increase subsequent Rab GTPase phosphorylation as part of a cooperative, feed-forward pathway.

LRRK2 is difficult to dye-label mono-molecularly, as the N-terminus is engaged in phosphoRab binding and the C-terminus is critical for activity. Nevertheless, analysis of the distribution of single molecule fluorescence intensity of our CF633-labeled LRRK2 preparation revealed a sharp peak, whether the preparation was evaluated immediately upon binding to Rab10 on bilayers (Figure 8–Figure Supplement 1A, B, D) or when spotted onto poly-lysine coated glass (Figure 8–Figure Supplement 1B, far right column). Panels A and B show the intensity at time t for large numbers of fluorescent molecules, either over 500 seconds (A) or 30 seconds (B). The intensity shift over time (panels A,B) may imply that the molecules slowly dimerize with a half time of 100-200 seconds, but additional work would be needed to confirm this. Continuous traces of the 30 longest lived spots showed that for some events this increase occurs even more quickly (1C). The fluorescent molecules remain on the bilayers for a significant period of time (panel E); moreover, when the molecules first bind to the surface, the single peak distribution of intensity does not change, irrespective of the time during the experiment that it actually binds (panel E). This gives us confidence that any changes observed were not occurring in solution and require Rab engagement. Note that we detect a minor species at log_2_=2.5 that constitutes between 2 and 6% of the molecules (panels D, F); this may represent dual labeled proteins and/or rare tetrameric complexes.

To confirm that LRRK2 Armadillo domain can bind both non-phosphorylated and phosphorylated Rabs simultaneously, GST-Rab8A was immobilized on glutathione agarose and Armadillo domain (1-552) protein pre-bound. Purified, phosphoRab10 was then added, and immunoblotting showed that phosphoRab10 bound to the beads only in the presence of Rab8A-anchored, Armadillo fragment (Figure 8 – Figure Supplement 2). Binding at both sites #1 and #2 is predicted to increase avidity of LRRK2 membrane association, consistent with our membrane recruitment data.

### PhosphoRab8 activates LRRK2 phosphorylation of Rab10 protein

The data presented thus far are consistent with apparent activation of LRRK2 by cooperative recruitment of the kinase to membrane microdomains enriched in Rab protein substrates. It was formally possible, however, that phosphoRab binding actually activates the kinase itself. To test this, we monitored the generation of phosphoRab10 using a highly specific monoclonal antibody in conjunction with immunoblotting. Rab10 protein was then phosphorylated by purified, full length LRRK2 kinase in vitro, with and without addition of pre-phosphorylated Rab8A protein. As shown in Figure 9 (A, C), the presence of stoichimetrically phosphorylated Rab8A (Dhekne et al., 2021) stimulated the rate of in vitro Rab10 phosphorylation by approximately 4 fold. Importantly, the ability of phosphoRab8A to stimulate LRRK2-mediated Rab10 phosphorylation required LRRK2’s K18 that is needed for phosphoRab binding (Fig. 9 B, D). We speculate that phosphoRab binding to the absolute N-terminus influences LRRK2’s higher order structure to stimulate kinase activity.

**Figure 9.**
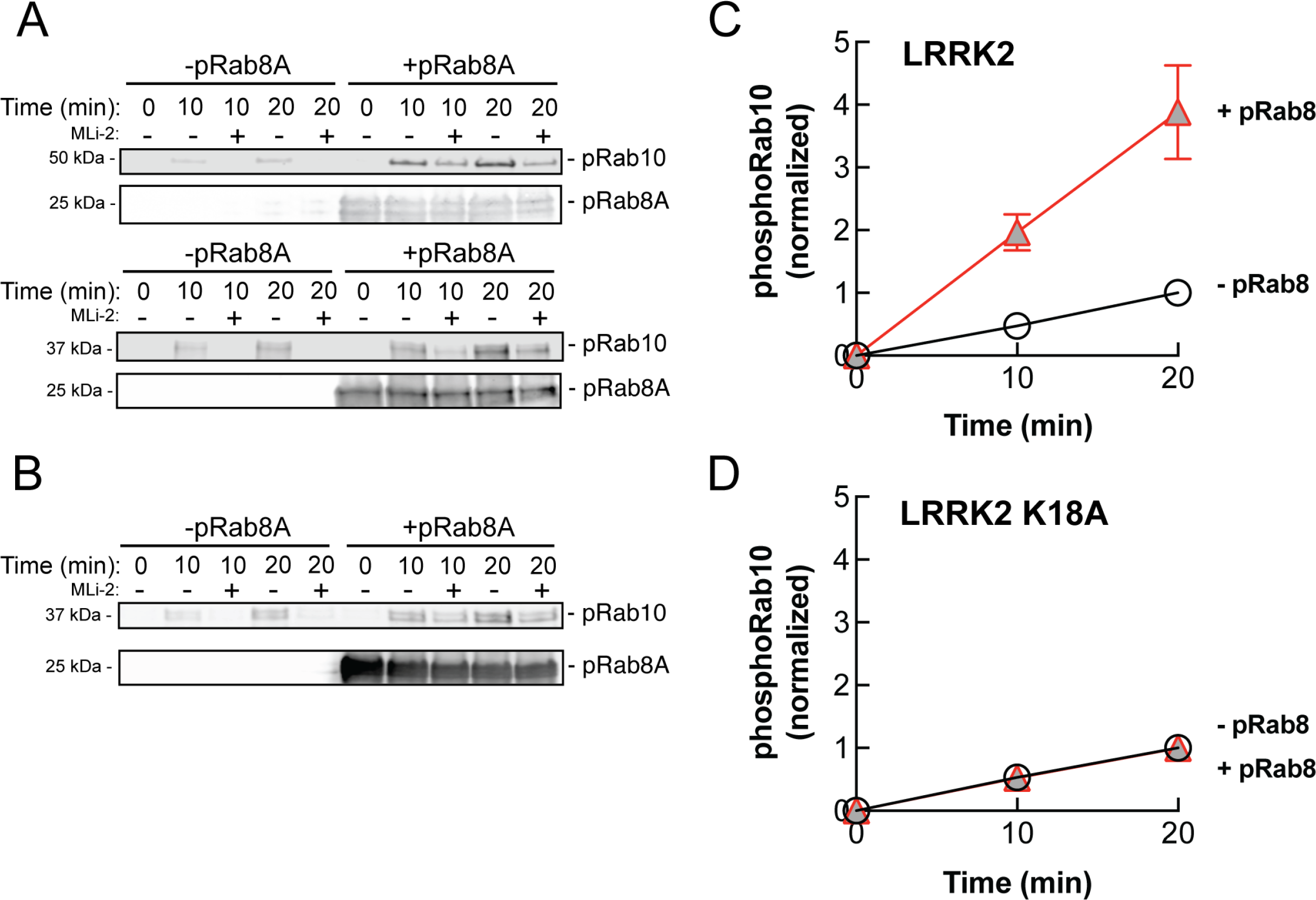
PhosphoRab8A activates LRRK2 phosphorylation of Rab10 in solution. A. Immunoblot analysis of the kinetics of LRRK2 G2019S phosphorylation of Rab10 with and without additional pRab8. Upper gel, GFP-Rab10 Q68L 1-181 substrate; Lower gel, His-Sumo-Rab10 wild type full length substrate. Indicated reactions contained 200nM MLi-2. pRab8A was detected with anti-phosphoRab8A antibody. B. Same as panel A with K18A-LRRK2-R1441G and His-Sumo-Rab10 wild type full length as substrate. PhosphoRab8A was detected with total Rab8 antibody. C. Kinetics of phosphoRab10 production as in A. Shown are the combined means of independent, quadruplicate determinations ± SEM, as indicated. D. PhosphoRab10 production as in B. Shown are the combined means of independent duplicate determinations, ±SEM, as indicated. Background signal in the presence of pRab8A is likely due to trace MST3 contamination that is not sensitive to MLi-2 inhibition and was subtracted. pRab8 preparation was by Method #1 for A, upper gel, and B, and Method #2 was used in panel A, lower gel.

## Discussion

LRRK2 is ∼90% cytosolic (Purlyte et al., 2019), and little was known about why membrane-associated LRRK2 appears to be much more active than the cytosolic pool of kinase. We have confirmed here that LRRK2 kinase relies upon substrate Rab GTPases to achieve membrane association, and revealed that LRRK2 utilizes two distinct Rab binding sites within its N-terminal Armadillo domain for this purpose. Site #1 (Fig. 5B) binds multiple, non-phosphorylated Rab substrates including Rab8A, Rab10 and Rab29, as well as the highly tissue specific- and non-substrate, Rab29-related, Rab32 and Rab38 proteins (Waschbüsch et al., 2014; McGrath et al., 2021). The second site (#2) is located at LRRK2’s absolute N-terminus, at a significant distance from the kinase active site; this site shows strong preference for phosphorylated Rab8A and Rab10 proteins. Our data show that both sites can be occupied simultaneously.

Figure 10 shows our current model for LRRK2 membrane recruitment. LRRK2 will interact reversibly with any one of the subset of Rab proteins that can bind to site #1. Rab29 shows the highest affinity for this site, but Rab8A can also bind with physiologically relevant affinity and is much more abundant in cells. Rab GTPases cluster in microdomains on distinct membrane surfaces (Pfeffer, 2017; Sönnichsen et al., 2000; DeRenzis et al., 2002; Barbero et al., 2002), thus this initial LRRK2 membrane association will bring the kinase in contact with other copies of the same substrate Rab proteins for phosphorylation. After an initial phosphorylation event, LRRK2 will then be held in place by bivalent association with one phosphorylated and one non-phosphorylated Rab protein. By binding to the kinase reaction product, LRRK2 enhances its effective, local activity by increasing the probability with which it will encounter another substrate Rab protein.

**Figure 10.**
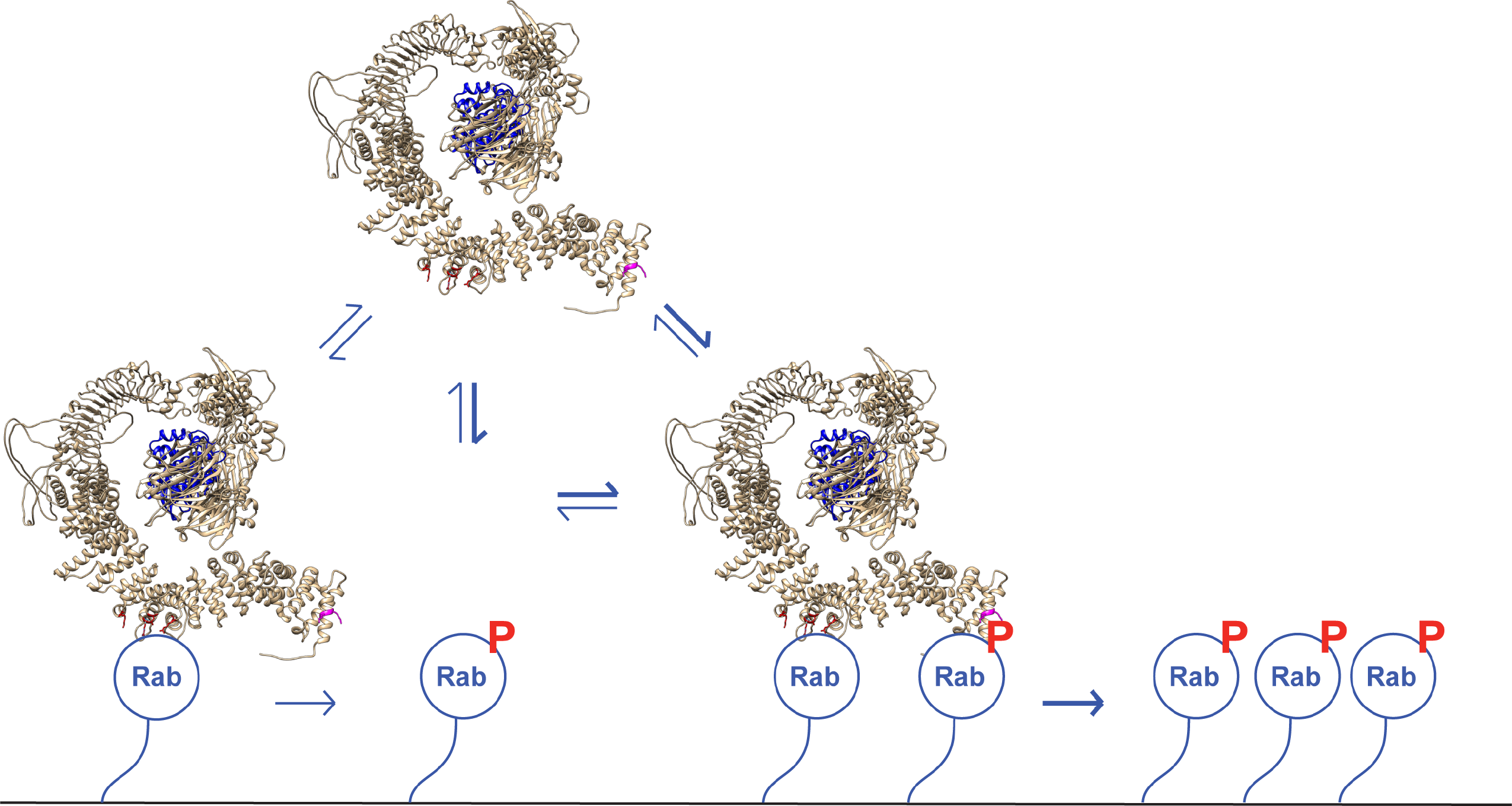
A model for LRRK2 membrane recruitment. LRRK2 can interact with non-phosphorylated Rab GTPases via site #1. Once membrane bound, it can generate phosphoRabs that can now engage site #2. Rab binding to both sites increases the avidity of LRRK2 for membranes and retains LRRK2 on the membrane surface to phosphorylate more Rab substrates. We have shown that LRRK2 binding to phosphoRabs also activates the kinase, likely by altering its oligomeric state.

Despite relatively similar affinities for their respective Rab binding partners, the phosphoRab-specific site appears to drive stable LRRK2 membrane association, as mutation of two key lysine residues strongly impacts co-localization of LRRK2 protein with phosphoRabs in cells. In addition, kinase activity leads to a much higher degree of LRRK2 association with planar lipid bilayers, despite the presence of binding Site #1 for non-phosphorylated-Rabs. Finally, K17/K18A LRRK2 that cannot bind to phosphorylated Rab proteins showed lower bilayer association in comparison with native LRRK2, confirming the importance of this interaction. LRRK2 phosphorylation of Rab GTPases is therefore required to form a new, additional interaction interface that greatly enhances the overall avidity of LRRK2 membrane association.

We also discovered that phosphoRab8A stimulates LRRK2 kinase action on Rab10 protein. We were not able to test the reverse scenario as the phosphoRab8A antibody is not adequately specific and cross-reacts with phosphoRab10 protein. Nevertheless, it seems very likely that phosphoRab10 will also activate LRRK2 for other substrate phosphorylation events. The most likely explanation is that phosphoRab binding to the LRRK2 N-terminus encourages an overall enzyme architecture that favors the active conformation. LRRK2 assumes multiple oligomeric states, and phosphoRab engagement and/or dual Rab engagement of the Armadillo domain likely influences the overall architecture of the enzyme.

It is important to note that quantitative mass spectrometry indicates that Rab10 is present at ∼600 times the copy number as LRRK2 in MEF cells and brain tissue (https://copica.proteo.info/#/copybrowse). Thus, if Rab10 is assumed to exist in cells at ∼2-5µM (Itzhak et al., 2016), LRRK2 will be present overall at about 3-8nM. These are very close to the concentrations used in our in vitro reconstitution experiments. Future experiments will be needed to elucidate the precise molecular state of LRRK2 upon engagement with Rab GTPases at sites #1 and #2.

Nichols et al. (2007) reported a single family with two affected siblings harboring LRRK2 E10K mutations. These patients presented with classic Parkinson’s disease symptoms at age 57 including bradykinesia, muscular rigidity, postural instability, and resting tremor. Compared with 46 G2019S LRRK2 patients in that study whose disease onset was on average, 63.5 years, the two siblings had a more severely disabling disease, as indicated by a higher Hoehn and Yahr assessment score (4 versus 2.5, where 5 represents confinement to bed or wheelchair unless aided). Our study provides a molecular explanation for how a mutation located far from the kinase or ROC-COR domains may cause Parkinson’s disease. We predict that the E10K mutation increases LRRK2 phosphoRab binding and membrane association and may display an even higher apparent activity than the most common pathogenic G2019S mutation. This distinction would need to be evaluated under conditions of MLi-2 washout, as exogenous expression would mask this subtle mechanistic feature.

The ability of multiple Rab binding sites to anchor LRRK2 on membranes will make the kinase appear more active than the pool of cytosolic LRRK2 protein. Rab binding may also increase access of LRRK2 to other kinases that stabilize it in a more active conformation. Anchoring LRRK2’s N-terminus may also influence autophosphorylation, which could also drive LRRK2 towards a more catalytically active conformation. Future structural studies of membrane anchored LRRK2 will provide important, additional information related to all of these possibilities.

## KEY RESOURCE TABLE

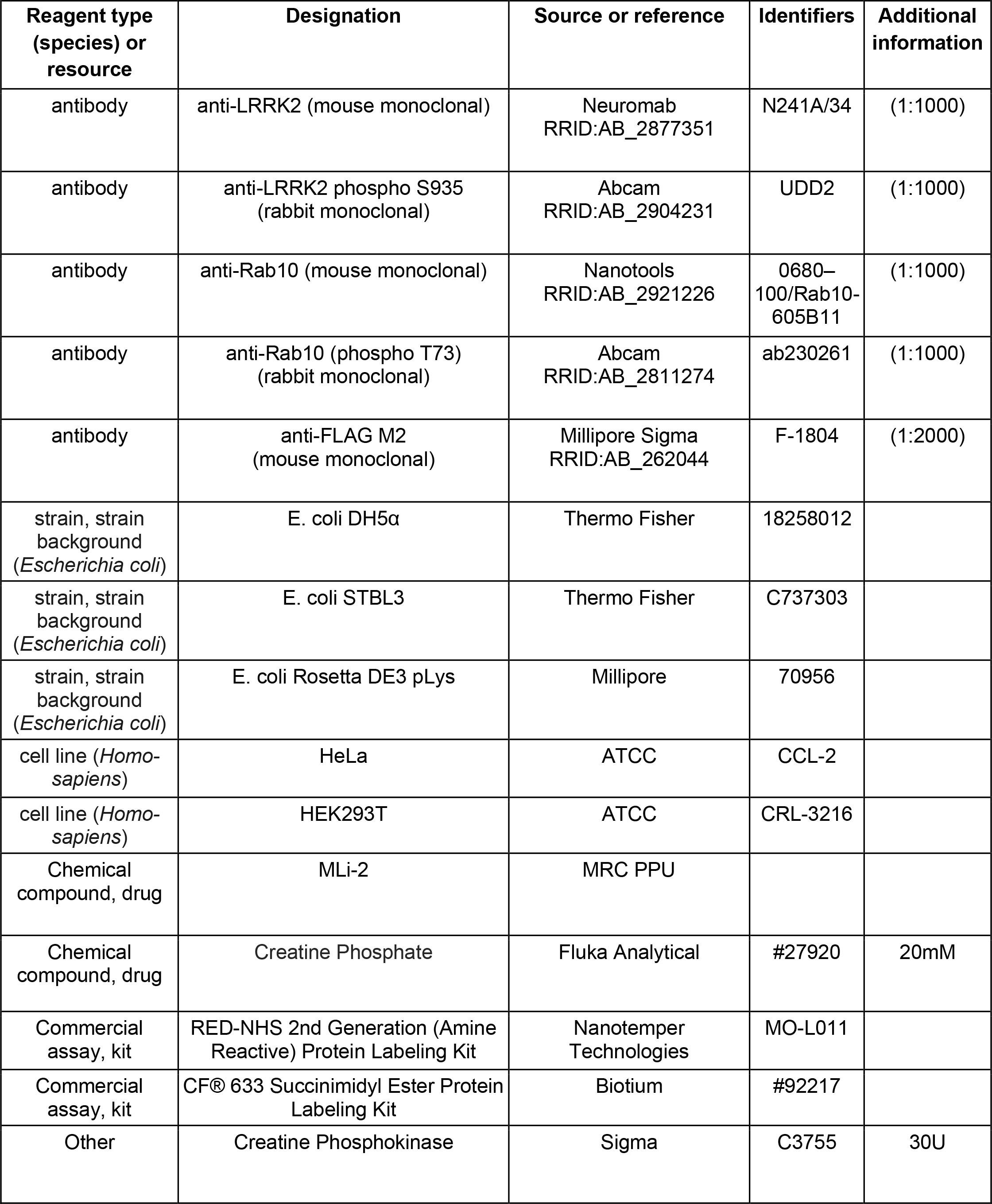
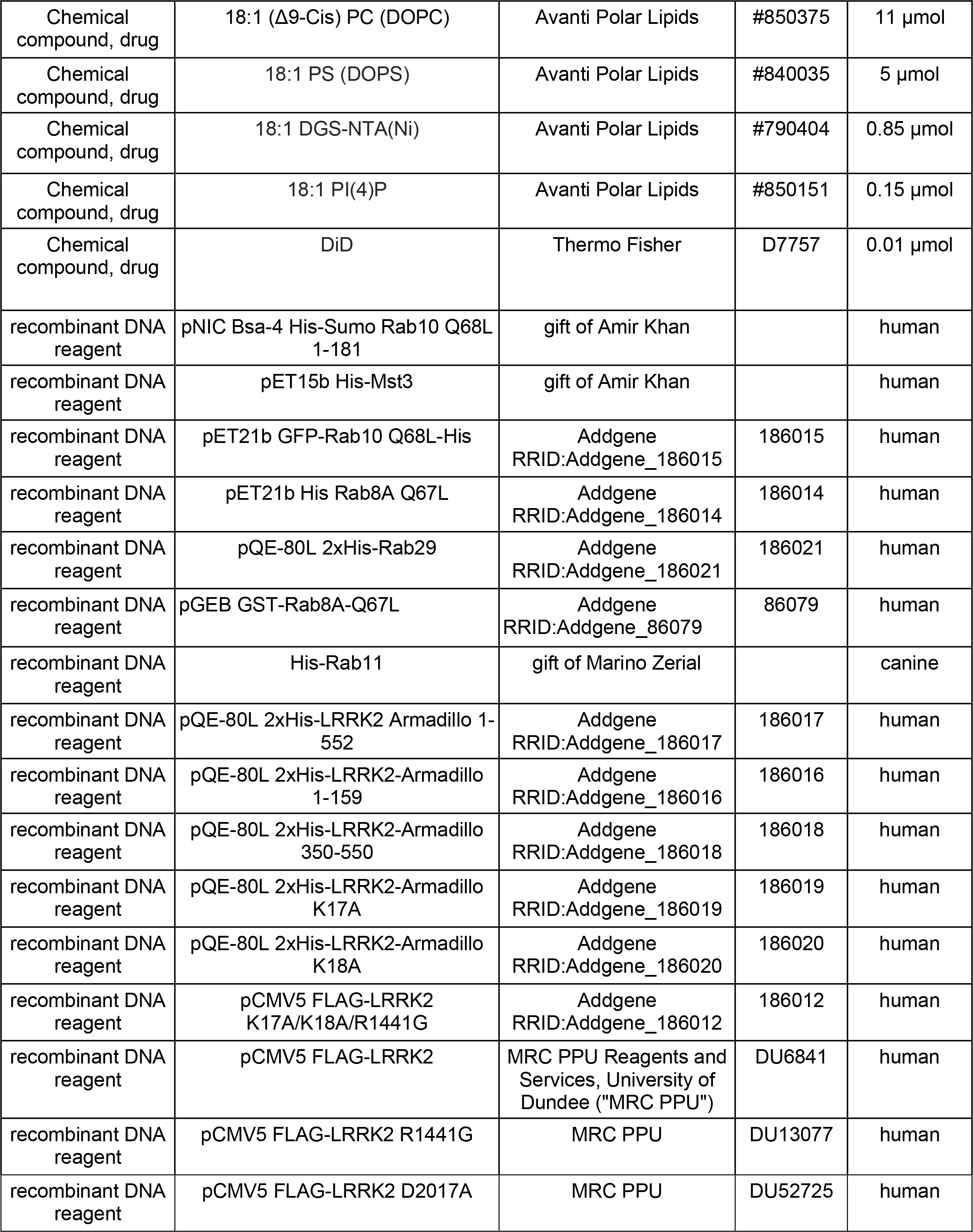
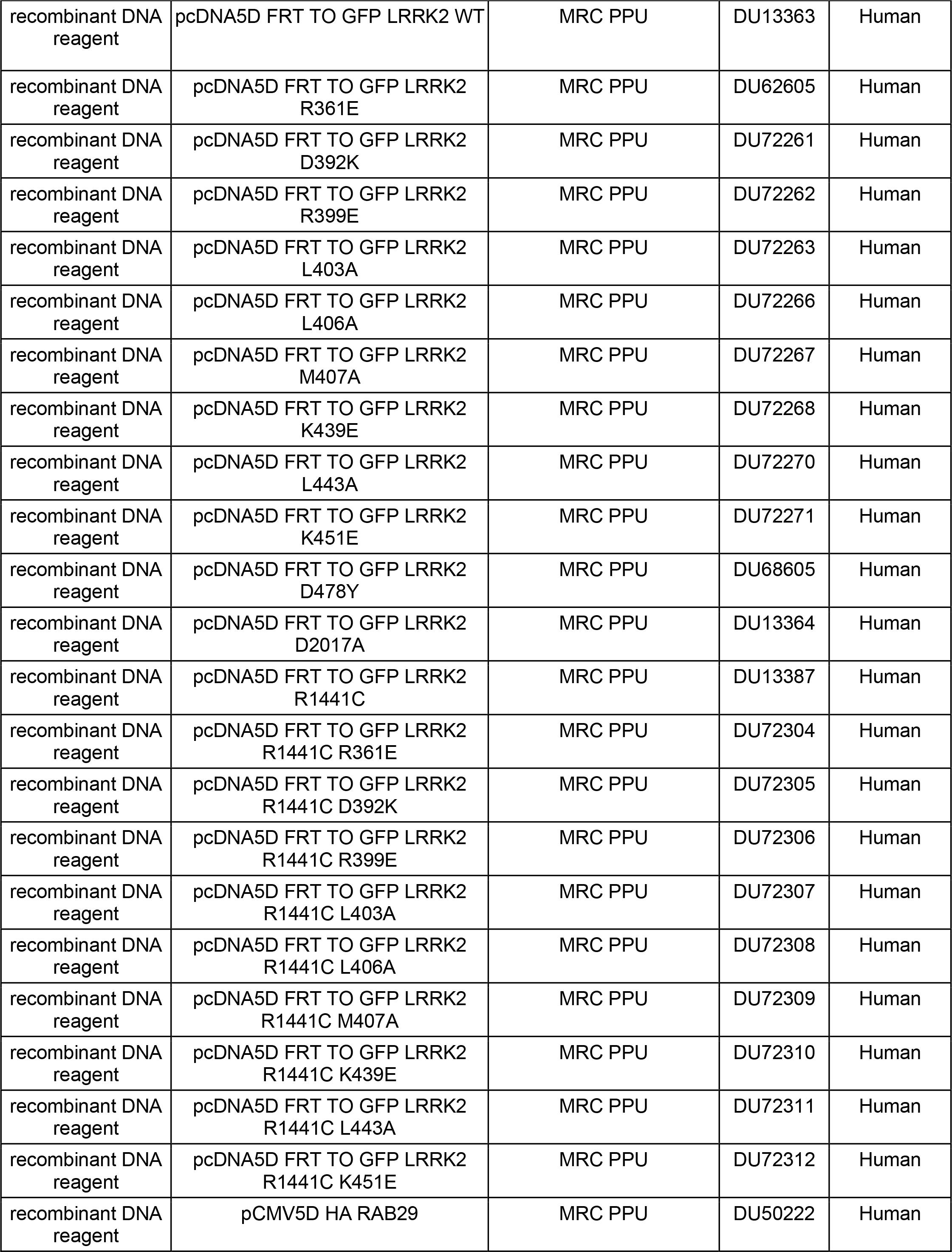
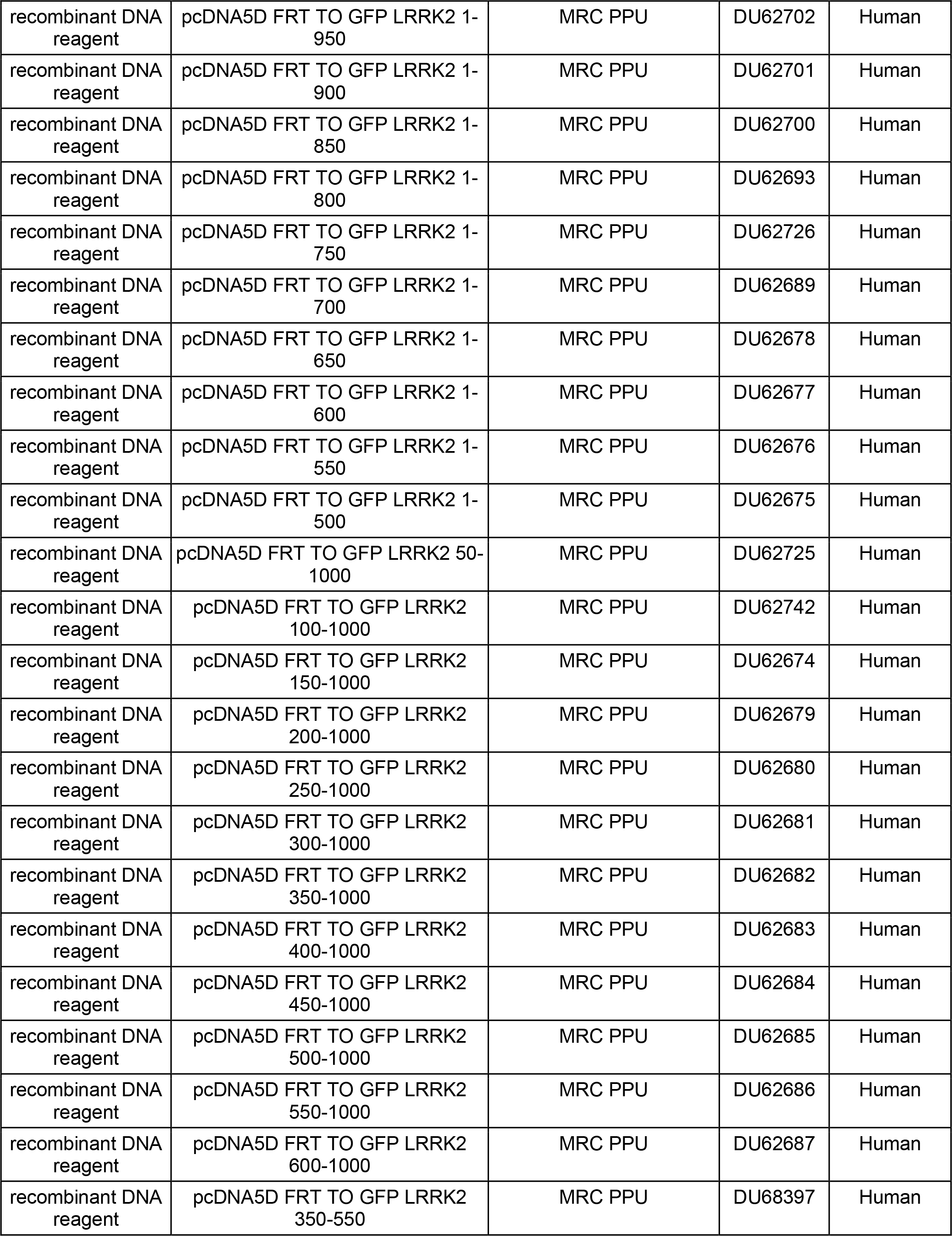
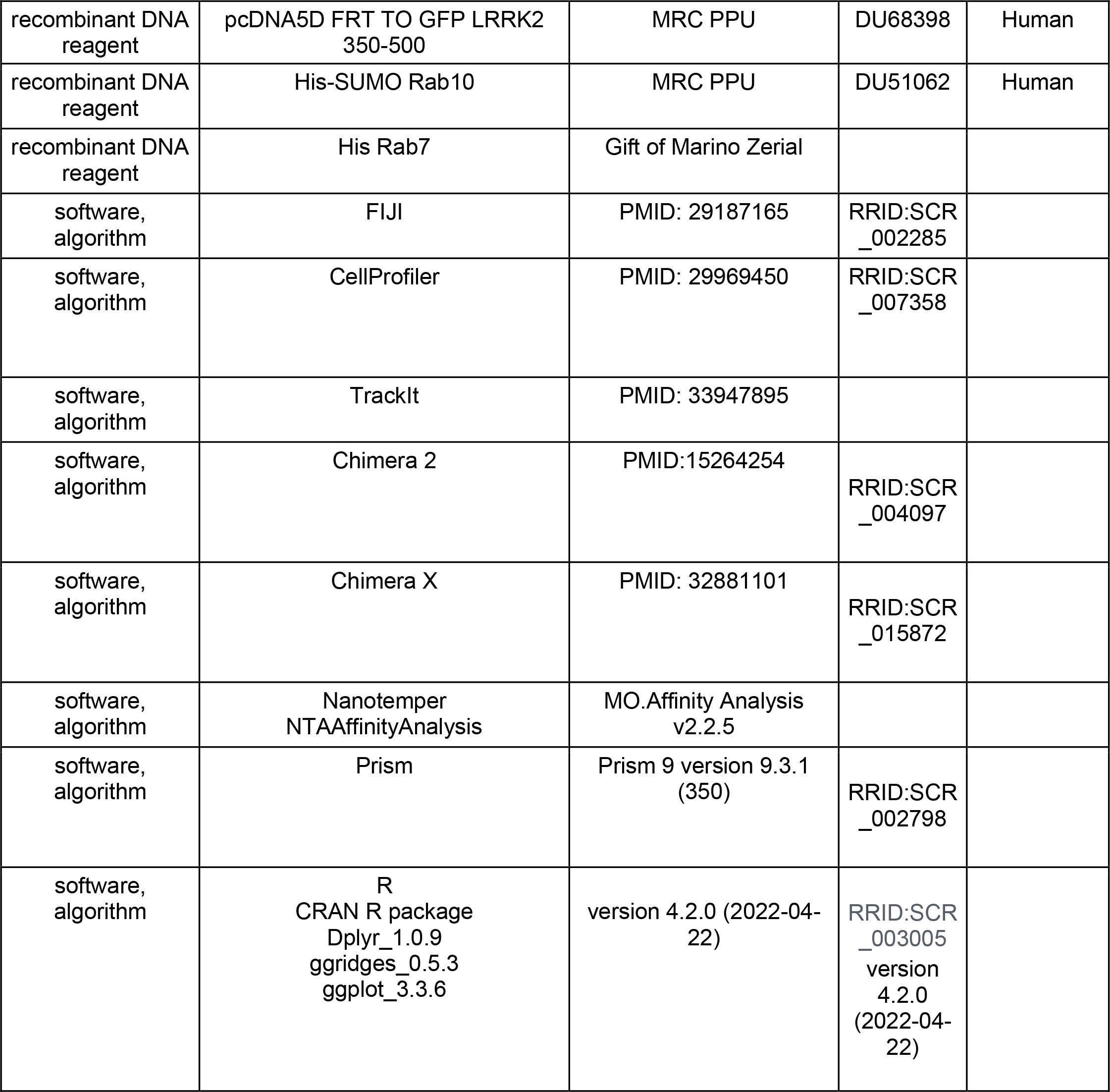

## Methods

### Cloning and plasmids

DNA constructs were amplified in Escherichia coli DH5α or STBL3 and purified using mini prep columns (Econospin). DNA sequence verification was performed by Sequetech (http://www.sequetech.com). pNIC Bsa-4 His-Sumo Rab10 Q68L 1-181 and pET15b His-Mst3 were kind gifts of Amir Khan (Harvard University). pET21b GFP-Rab10 Q68L-His was subcloned from GFP-Rab10 (Gomez et al. 2019) into pET21b. The C-terminal His tagged version was generated by Gibson assembly. His Rab8A Q67L was subcloned from HA-Rab8A (DU35414, Medical Research Council at Dundee) into pET14b. Point mutations were generated using site directed mutagenesis. His-Rab29 wild type was subcloned from HA-Rab29 (DU5022, Medical Research Council at Dundee) into the pQE-80L backbone. pCMV5 FLAG-LRRK2 (DU6841), Flag-LRRK2 R1441G (DU13077), His-SUMO Rab10 (DU51062) and FLAG-LRRK2 D2017A (DU52725) were obtained from the Medical Research Council at Dundee. His-Armadillo 1-552, 1-159, and 350-550 were all cloned from pCMV5 FLAG-LRRK2 into pQE-80L. K17A, K18A, and K17A/K18A LRRK2 and LRRK2 Armadillo were generated using site directed mutagenesis. All cloning and subcloning was done by Gibson assembly.

### Rab GTPase, LRRK2 Armadillo domain, and LRRK2 purification

His Rab29, His Rab10-Q68L (1–181), His-Mst3, His-Rab8A Q67L, His-LRRK2 Armadillo (1-552), His-LRRK2 Armadillo (1-159), His-LRRK2 Armadillo (350-550), His-LRRK2 Armadillo K17A, His-LRRK2 Armadillo K18A, and GST-Rab8A Q67L were purified in E.coli BL21 (DE3 pLys). Detailed protocols can be found in Gomez et al., 2020 https://dx.doi.org/10.17504/protocols.io.bffrjjm6 and Vides et al., 2021 https://dx.doi.org/10.17504/protocols.io.bvvmn646. Bacterial cells were grown at 37°C in Luria Broth and induced at A600 nm = 0.6–0.7 by the addition of 0.3 mM isopropyl-1-thio-β-D-galactopyranoside (Gold Biotechnology) and harvested after 18 h at 18°C. The cell pellets were resuspended in ice cold lysis buffer (50 mM HEPES, pH 8.0, 10% [vol/vol] glycerol, 500 mM NaCl,10 mM imidazole (for His-tagged purification only), 5 mM MgCl_2_, 0.2mM tris(2-carboxyethyl) phosphine (TCEP), 20 μM GTP, and EDTA-free protease inhibitor cocktail (Roche). The resuspended bacteria were lysed by one passage through an Emulsiflex-C5 apparatus (Avestin) at 10,000 lbs/in^2^ and centrifuged at 40,000 rpm for 45 min at 4°C in a Beckman Ti45 rotor. Cleared lysate was filtered through a 0.2µm filter (Nalgene) and passed over a HiTrap TALON crude 1mL column (Cytiva) for His-tagged proteins or a GSTrap High Performance 1 mL column (Cytiva) for GST-tagged proteins. The column was washed with lysis buffer until absorbance values reached pre-lysate values. Protein was eluted with a gradient from 20-500 mM imidazole containing lysis buffer for His-tagged proteins or 0-50 mM reduced glutathione containing lysis buffer for GST-tagged proteins. Peak fractions analyzed by 10% SDS-PAGE to locate protein. The eluate was buffer exchanged and further purified by gel filtration on Superdex-75 (GE Healthcare) with 50 mM HEPES, pH 8, 5% (vol/vol) glycerol, 150 mM NaCl, 5 mM MgCl_2_, 0.1 mM tris(2-carboxyethyl) phosphine (TCEP), and 20 μM GTP.

LRRK2 R1441G was transfected into HEK293T cells with Polyethylenimine HCl MAX 4000 (PEI) (Polysciences, Inc.) and purified 48hrs post transfection. Cells were lysed in 50mM Hepes pH 8, 150mM NaCl, 1mM EDTA, 0.5% Triton-X 100, 10% (vol/vol) glycerol and protease inhibitor cocktail (Roche). Lysate was centrifuged at 15,000g for 20min in Fiberlite F15 rotor (ThermoFischer). Clarified lysate was filtered through 0.2µm syringe filters and circulated over anti-FLAG M2 affinity gel (Sigma) at 4°C for 4hrs using a peristaltic pump. The affinity gel was washed with 6 column volumes of lysis buffer followed by 6 column volumes of elution buffer (50mM Hepes pH 8, 150mM NaCl, and 10% [vol/vol] glycerol). Protein was eluted from resin with 5 column volumes of FLAG peptide (0.25mg/mL) containing elution buffer. Eluate was supplemented to 20µM GTP, 1mM ATP, and 2mM MgCl_2._

### In vitro Rab phosphorylation and microscale thermophoresis

A detailed method can be found at https://dx.doi.org/10.17504/protocols.io.bvvmn646). His-Rab10 Q68L 1–181 or His-Rab8A Q67L was incubated with His-Mst3 kinase at a molar ratio of 3:1 (substrate:kinase). The reaction buffer was 50 mM Hepes, pH 8, 5% (vol/vol) glycerol, 100 mM NaCl, 5 mM MgCl_2_, 0.2 mM TCEP, 20 µM GTP, 5 μM BSA, 0.01% Tween-20, and 2 mM ATP (no ATP for negative control). The reaction mixture was incubated at 27°C for 30 min in a water bath. Phosphorylation completion was assessed by Western blot. Immediately after phosphorylation, the samples were transferred to ice before binding determination. See also https://dx.doi.org/10.17504/protocols.io.bvjxn4pn.

Protein–protein interactions were monitored by microscale thermophoresis using a Monolith NT.115 instrument (Nanotemper Technologies). His LRRK2 Armadillo (1-552), (1-159), (350-550), K17A and K18A were labeled using RED-NHS 2^nd^ Generation (Amine Reactive) Protein Labeling Kit (Nanotemper Technologies). For all experiments, the unlabeled protein partner was titrated against a fixed concentration of the fluorescently labeled LRRK2 Armadillo (100 nM); 16 serially diluted titrations of the unlabeled protein partner were prepared to generate one complete binding isotherm. Binding was carried out in a reaction buffer in 0.5-ml Protein LoBind tubes (Eppendorf) and allowed to incubate in the dark for 30 min before loading into NT.115 premium treated capillaries (Nanotemper Technologies). A red LED at 30% excitation power (red filter, excitation 605–645 nm, emission 680–685 nm) and IR-laser power at 60% was used for 30 s followed by 1 s of cooling. Data analysis was performed with NTAffinityAnalysis software (NanoTemper Technologies) in which the binding isotherms were derived from the raw fluorescence data and then fitted with both NanoTemper software and GraphPad Prism to determine the *K*_D_ using a non-linear regression method. The binding affinities determined by the two methods were similar. Shown are averaged curves of Rab GTPase binding partners from single readings from two different protein preparations. Note that the affinities reported here are underestimates as preps of His Rab10-Q68L (1–181) and His-Rab8A Q67L routinely contained a 50:50 ratio of bound GTP:GDP as determined by mass spectroscopy; data were not corrected for this.

### Cell culture and Immunoblotting

HEK293T and HeLa cells were obtained from American Type Culture Collection were cultured at 37°C and under 5% CO2 in Dulbecco’s modified Eagle ’s medium containing 10% fetal bovine serum, 2 mM Glutamine, and penicillin (100 U/ml)/streptomycin (100 μg/ml). HEK293T and HeLa cells were transfected with polyethylenimine HCl MAX 4000 (Polysciences). Cells were routinely checked for Mycoplasma by PCR analysis.

HeLa cells for pRab10 recovery kinetics were lysed 48 hr post transfection and MLi-2 treatment in ice cold lysis buffer (50mM Tris pH 7.4, 150mM NaCl, 0.5% Triton X-100, 5mM MgCl_2_, 1 mM sodium orthovanadate, 50 mM NaF, 10 mM 2-glycerophosphate, 5 mM sodium pyrophosphate, 0.1 μg/ml mycrocystin-LR (Enzo Life Sciences), and EDTA-free protease inhibitor cocktail (Sigma-Aldrich). Lysates were centrifuged at 14,000g for 15 min at 4°C and supernatant protein concentrations were determined by Bradford assay (Bio-Rad).

A detailed protocol for blotting is available on protocols.io (Tonelli and Alessi, https://dx.doi.org/10.17504/protocols.io.bsgrnbv6). 20µg of protein were run on SDS PAGE gels and transferred onto nitrocellulose membranes using a Bio-Rad Trans-turbo blot system. Membranes were blocked with 2% BSA in Tris-buffered saline with Tween-20 for 30 at RT. Primary antibodies used were diluted in blocking buffer as follows: mouse anti-LRRK2 N241A/34 (1:1000, Neuromab); rabbit anti-LRRK2 phospho S935 (1:1000, Abcam); mouse anti-Rab10 (1:1000, Nanotools); and rabbit anti-phospho Rab10 (1:1000, Abcam). Primary antibody incubations were done overnight at 4°C. LI-COR secondary antibodies diluted in blocking buffer were 680 nm donkey anti-rabbit (1:5,000) and 800 nm donkey anti-mouse (1:5,000). Secondary antibody incubations were for 1 h at RT. Blots were imaged using an Odyssey Infrared scanner (LI-COR) and quantified using ImageJ software.

### MLi-2 washout/ pRab10 recovery kinetics

As described by Ito et al. (2016), HeLa cell seeded in 6x 60mm dishes expressing FLAG-LRRK2, LRRK2 K17A/K18A, LRRK2 R1441G, or LRRK2 R1441G/K17A/K18A for 48 hours were incubated with 200nm MLi-2 or DMSO for 1 hour under normal growth conditions at 37°C. To remove the MLi-2 inhibitor, cells were washed 4 times with complete media. Washouts were done to allow for 120-15 min of enzyme activity recovery; after which, cells were harvested.

### Confocal light microscopy

The standard method to obtain images in Figure 3 and Figure 3 supplements can be found on protocols.io (Purlyte et al., 2022 https://dx.doi.org/10.17504/protocols.io.b5jhq4j6). For Figure 8, cells were plated onto collagen coated coverslips with indicated plasmids. Cells were washed with ice cold phosphate buffered saline (PBS) 3x. After, they were incubated in glutamate buffer (25mM KCl, 25mM Hepes pH7.4, 2.5mM magnesium acetate, 5mM EGTA, and 150mM K glutamate) for 5 min on ice. Coverslips were dipped into liquid nitrogen and held for 5 sec before removal. They were thawed at RT, incubated in glutamate buffer for 2 min and then in PBS for 5 min. Cells were fixed with 3.5% paraformaldehyde in PBS for 15 min, permeabilized for 3 min in 0.1% Triton X-100, and blocked with 1% BSA in PBS. Antibodies were diluted as follows: mouse anti-FLAG (1:2000, Sigma-Aldrich) and rabbit anti pRab10 (1:2000; Abcam). Highly cross-absorbed H+L secondary anti-bodies (Life Technologies) conjugated to Alexa Fluor 568 or 647 were used at 1:2,000. Images were obtained using a spinning disk confocal microscope (Yokogawa) with an electron multiplying charge coupled device camera (Andor) and a 100× 1.4 NA oil immersion objective. Mander’s correlation coefficients were calculated by analyzing maximum intensity projection images with CellProfiler software (Carpenter et al., 2006; Stirling et al., 2021).

Co-localization of Rab29 with full length LRRK2 and its mutants was quantified using an unbiased Cellprofiler pipeline as follows: Step 1. Imported raw .lsm files; 2. metadata extracted from the file headers; 3. images grouped by mutations and split into 3 channels; 4. nuclei identified as primary objects after rescaling intensities; 5. Nucleus is defined as the primary object and cells are identified by ‘propagation’ as secondary objects; cells are identified as the using the rescaled and smoothened LRRK2 channel. 6. Co-localization within whole cells is measured by thresholded (10) Mander’s coefficient on the entire batch of images. Data plotted from Cellprofiler are relative values.

### Substrate supported lipid bilayer preparation

A detailed method can be found at <u>dx.doi.org/10.17504/protocols.io.x54v9y7qzg3e/v1</u>. Briefly, we used Lab-TeKII 8 chambered No. 1.5 borosilicate cover glasses (Fischer) for LRRK2 recruitment assays. Reactions chambers were cleaned by 30 min incubation in Piranha solution (1:3 [vol/vol] ratio of 30% H_2_O_2_ and 98% H_2_SO_4_) and extensive washing in Milli-Q water. The reaction chambers were stored in Milli-Q water for up to 2 weeks. Before use, reaction chambers were dried and further cleaned in a Harrick Plasma PDC-32C plasma cleaner for 10 min at 18W under ambient air.

We prepared substrate supported lipid bilayers on glass coverslips with 65% DOPC, 29% DOPS, 5% DOGS-NTA[Ni^2^], 1% PI(4)P, 0.01% DIL (Avanti Polar Lipids;Thermo). The lipid mixture was suspended in 1ml chloroform and then dried under nitrogen flow in a glass vial and kept under vacuum for at least 1 h. The dried lipids were hydrated in SLB buffer (20 mM Hepes pH 8, 150 mM potassium acetate, 1 mM MgCl_2_) by vortexing to produce multilamellar vesicles (MLVs). SUVs were prepared by bath sonication followed by extrusion through 100 nm polycarbonate membrane 21 times (Avestin). The produced SUVs were stored at -20°C. The supported lipid bilayer was formed in cleaned reaction chambers on glass surfaces by addition of liposomes to a final concentration of 5mM liposomes in SLB buffer. SUV fusion was induced by addition of 1mM CaCl_2_ and incubated for 45 min at 37°C. Next, the unfused vesicles were washed with Milli-Q water and STD buffer (20mM Hepes pH 8, 150mM NaCl, 5mM MgCl_2_).

Lab-TeKII 8 chambered No. 1.5 borosilicate coverglass (Fisher) were coated with Poly-D-lysine as follows (Adhikari et al., 2022). 10 mg Poly-D-lysine (MPBio # SKU:02150175-CF) was dissolved in 1ml of sterile Milli-Q water as a 1% stock solution. The stock solution was then diluted two fold in PBS as 1X coating solution. Coating solution (200µl) was added to the reaction chamber and incubated for 5 min at 37°C. The coating solution was then removed by rinsing the chamber thoroughly with sterile Milli-Q water and equilibrated with reaction buffer (20mM Hepes pH8; 150mM NaCl, 5mM MgCl2, 4mM ATP, 20µM GTP, 20 mM creatine phosphate, 30U creatine phosphokinase) (<u>dx.doi.org/10.17504/protocols.io.x54v9y7qzg3e/v1</u>).

### TIRF microscopy

A detailed method can be found on protocols.io (Adhikari et al., 2022 <u>dx.doi.org/10.17504/protocols.io.x54v9y7qzg3e/v1</u>). All LRRK2 recruitment movies were obtained at 25°C, at a frame rate capture interval of 1 sec using a Nikon Ti-E inverted microscope with the Andor iXon+EMCCD camera model DU885 with PerfectFocus and a Nikon TIRF Apo 100x 1.46 NA oil immersion objective. The imaging was done with 300 EM camera gain and 50 ms exposure time with 200µW laser intensity. We analyzed the microscopy data with TrackIt (Kuhn T., et al., 2021) to obtain spot density of bound LRRK2.

### Rab10-dependent LRRK2 recruitment

A detailed method can be found on protocols.io (Adhikari et al., 2022 <u>dx.doi.org/10.17504/protocols.io.x54v9y7qzg3e/v1</u>). Purified FLAG LRRK2 is labeled with CF633 succinimidyl ester (Biotium 92217) by incubation with dye for 1 hr at RT in the dark. After dye removal using Pierce Dye Removal Columns (Thermo Scientific #22858), protein was determined by Bradford Assay. Labeling efficiency was determined using the dye extinction coefficient and preps were labeled with 2-3 moles of dye per mole LRRK2 for all experiments.

GFP Rab10 Q68L C-terminal His was added to supported lipid bilayers at a final concentration of 2.5µM in STD buffer and incubated for 20 min at 37°C. After incubation, Rab coated supported lipid bilayers were washed with STD buffer and then equilibrated with reaction buffer (20mM Hepes pH 8, 150mM NaCl, 5mM MgCl_2_, 4mM ATP, 20μM GTP, 20mM creatine phosphate, 30U creatine phosphokinase). 14nM CF633-FLAG LRRK2 was prepared in reaction buffer and allowed to equilibrate to RT for 5 min. 40 sec into imaging, 100µL from the 200uL in the reaction chamber was removed. At 60 sec, 100µL of 14nM CF FLAG LRRK2 was added and imaged for 600 sec and for 300 sec for no Rab10 control.

### LRRK2 Kinase Activation Assay

Method #1. Purified His-Rab8A Q67L (0.5 mg) was phosphorylated using His-MST3 kinase (0.1-0.3 mg) as described above at 30°C overnight in MST3 reaction buffer (50 mM HEPES, pH 8, 5% (v/v) glycerol, 150 mM NaCl, 5 mM MgCl_2_, 0.2 mM TCEP, 20 µM GTP, 5 µM BSA, 0.01% Tween-20, and 2 mM ATP). Phosphorylated Rab8A (25 kDa) was then resolved from MST3 (55 kDa) by gel filtration on a 24 mL Superdex 75 10/300 column (Cytiva Life Sciences, #17517401. An additional Method #2 was attempted to try to further remove trace MST3 from phosphoRab8. GST-PreScission protease was bound to glutathione agarose. His-MST3 was added to the beads and incubated overnight at 4°C. The supernatant containing free MST3 was then passed through a Nickel-NTA column to remove any uncleaved His-MST3. The pooled, untagged, MST3 supernatants were then used to phosphorylate His-Rab8A. The products of this reaction were gel filtered on Superdex 75 column as before, and phosphorylated His-Rab8A was then further purified by immobilization on nickel-NTA agarose, eluted with 500 mM imidazole after washing, and desalted as described above.

LRRK2 G2019S (88 nM; Thermo Fisher Scientific #A15200) or purified FLAG-LRRK2 R1441G K18A (Adhikari et al., 2022) was incubated with 3 µM His-GFP-Rab10 Q68L (1-181) or His-SUMO-Rab10 wild type full length substrate ± 6 µM phosphorylated Rab8A Q67L in 50 mM HEPES pH 8, 5% (v/v) glycerol, 150 mM NaCl, 10 mM MgCl_2_, 250 µM GTP, 5 µM BSA, and 2 mM ATP. No difference was detected between the two Rab10 substrates. The reaction was incubated at 30°C in a water bath. Reactions were stopped by the addition of SDS-PAGE sample buffer; MLi-2 (200nM) was added to control reactions. Samples were analyzed by SDS-PAGE and immunoblotted for phosphoRab10. Blots were imaged using Li-COR and bands quantified using ImageJ. The values obtained with MLi-2 were subtracted from their respective timepoints to monitor LRRK2-dependent phosphorylation; background was due to trace residual MST kinase. Values from four independent, replicate experiments were normalized to the 20 min time point and plotted together using GraphPad Prism.

### Dual Rab GTPase Binding to the LRRK2 Armadillo Domain

The strategy was to immobilize Rab8A, bind Armadillo domain, and then test if Rab8A-tethered Armadillo domain could simultaneously bind phosphoRab10. His-Rab10 Q68L 1-181 was pre-phosphorylated with His-MST3 kinase at a molar ratio of 3:1 (substrate:kinase) at 30°C for 2 hours in MST3 reaction buffer. 50 µL glutathione agarose slurry was pelleted and resuspended in 50 mM HEPES, pH 8, 5% (v/v) glycerol, 150 mM NaCl, 5 mM MgCl_2_, 0.2 mM TCEP, 100 µM GTP, 5 µM BSA, 0.01% Tween-20 to achieve a total volume of 50 µL. GST-Rab8A Q67L (6µM in 50µl) was incubated with glutathione beads in reaction buffer for 30 min at room temperature (RT) on a rotator. The reaction was spun down at 3200 x g for 30 sec and the supernatant discarded. His-LRRK2 Armadillo domain 1-552 in reaction buffer (or buffer alone) was added to beads to achieve a final concentration of 10 µM in 50 µL and incubated for 30 min at RT on a rotator. The reaction was spun down as before and the supernatant discarded. Phosphorylated His-Rab10 Q68L 1-181 (4µM final) were added to beads in a final volume of 50 µl. Reactions were incubated for 30 min at RT on a rotator. The reaction was spun down at 3200 xg for 30 seconds and the supernatant discarded; reaction buffer (500 µl) was used to wash the beads twice. Proteins were eluted from the beads using 50 µl elution buffer (50 mM HEPES, pH 8, 5% (v/v) glycerol, 150 mM NaCl, 5 mM MgCl_2_, 0.2 mM TCEP, 20 µM GTP, 50 mM reduced glutathione). The reaction was spun down at 3200 xg for 30 sec and the supernatant was collected. Samples were then analyzed by SDS-PAGE and immunoblotted for phosphoRab10. Blots were imaged using Li-COR, and bands were quantified using ImageJ.

### Intensity analysis of TIRF videos

Tracks of individual molecules were extracted from TIRF microscopy images using the TrackIT FiJi plugin and converted to .csv files using the custom “getTracks.m” Matlab script (https://github.com/PfefferLab/Vides_et_al_2022). These files were loaded as data frames in R and processed with dplyr for the binning and normalization steps. Pre-normalized intensities *I_t_* were obtained from the amplitude value fitted by TrackIT (backround corrected amplitude of the Gaussian fit of each particle). Ridge plots were produced using the ggridges package with a Gaussian Kernel density and a bandwidth of 0.2. Code used to generate each figure is available on GitHub (https://github.com/PfefferLab/Vides_et_al_2022).

## Acknowledgements

This study was funded by the joint efforts of The Michael J. Fox Foundation for Parkinson’s Research (MJFF) [17298 & 6986 (S.R.P. & D.R.A.)] and Aligning Science Across Parkinson’s (ASAP) initiative. MJFF administers the grant (ASAP-000463, S.R.P. & D.R.A.) on behalf of ASAP and itself. Funds were also provided by the Medical Research Council [grant no. MC_UU_00018/1 (D.R.A.), the pharmaceutical companies supporting the Division of Signal Transduction Therapy Unit (Boehringer-Ingelheim, GlaxoSmithKline, Merck KGaA (D.R.A.), and a Ph.D. fellowship from Consejería de Economía, Conocimiento y Empleo del Gobierno de Canarias in partnership with Fondo Social Europeo (E.S-L.).

For the purpose of open access, the authors have applied a CC-BY public copyright license to the Author Accepted Manuscript version arising from this submission. All primary data associated with each figure has been deposited in a repository and can be found at https://doi.org/10.5061/dryad.3tx95x6j7; quantitation data of the blots in Figure 3--Fig. Supp. 4 (for the bar graphs in Figures 3C and 3D) can be found at doi (10.5281/zenodo.7057419).

We are especially grateful to Dr. Gheorghe Chistol for helpful discussion and Chloe Rollock for help with protein purification. We also thank the excellent technical support of the MRC protein phosphorylation and ubiquitylation unit (PPU) DNA sequencing service (coordinated by Gary Hunter), the MRC-PPU tissue culture team (coordinated by Edwin Allen), MRC-PPU Reagents and Services antibody and protein purification teams (coordinated by Dr James Hastie).

---

**Figure 3 – Figure Supplement 1.**
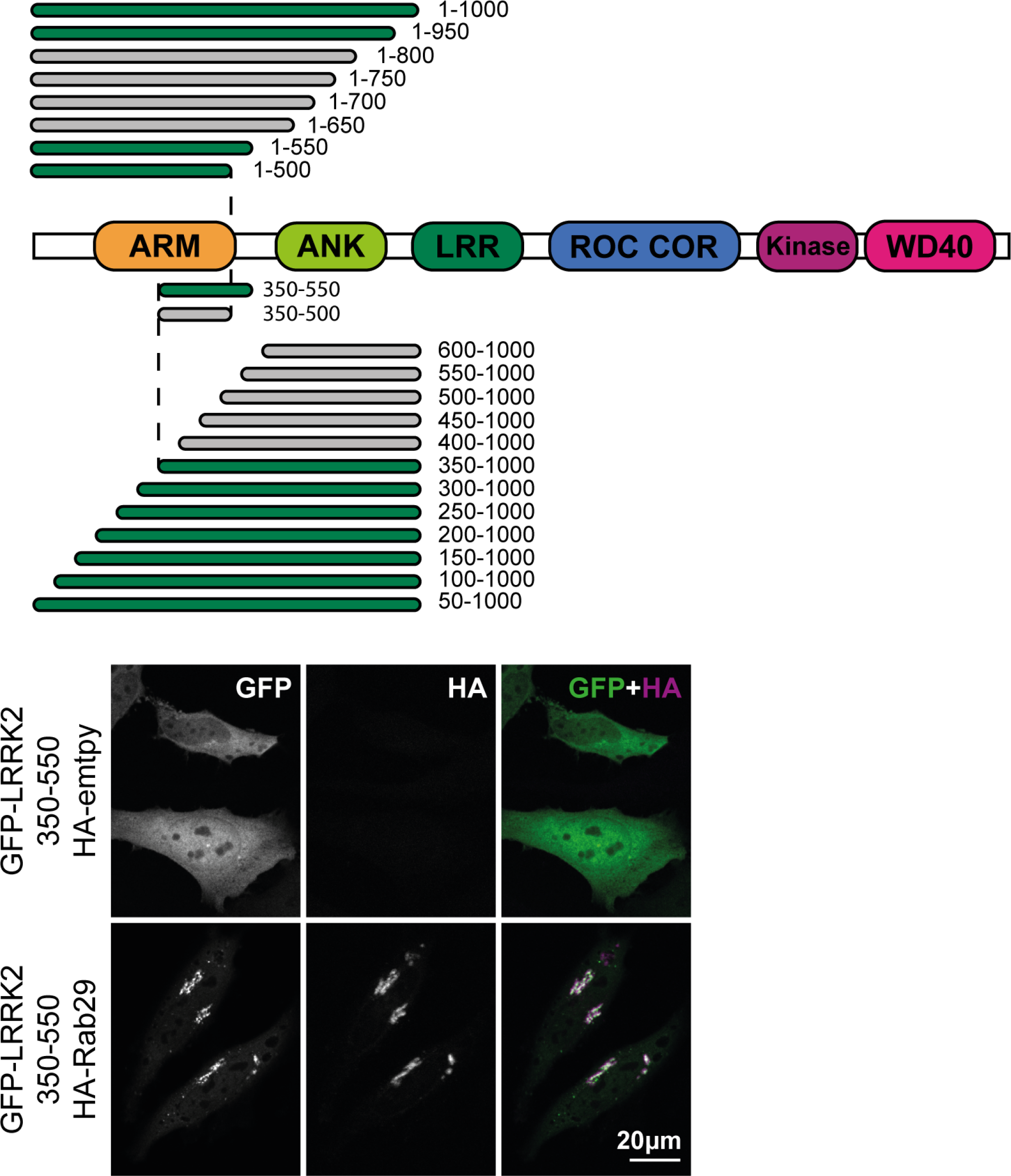
(top) Fragments of GFP-LRRK2 that were co-expressed with HA-Rab29 in HeLa cells. 24h post transfection, cells were fixed and localization assessed by confocal microscopy. Fragments that co-localized with Rab29 at the Golgi are shown in green and those that failed to co-localize in gray. The smallest fragment of LRRK2 that co-localized with Rab29 encompassed residues 350-550 (shown below). Magnification bar, 20µm.

**Figure 3 – Figure Supplement 2.**
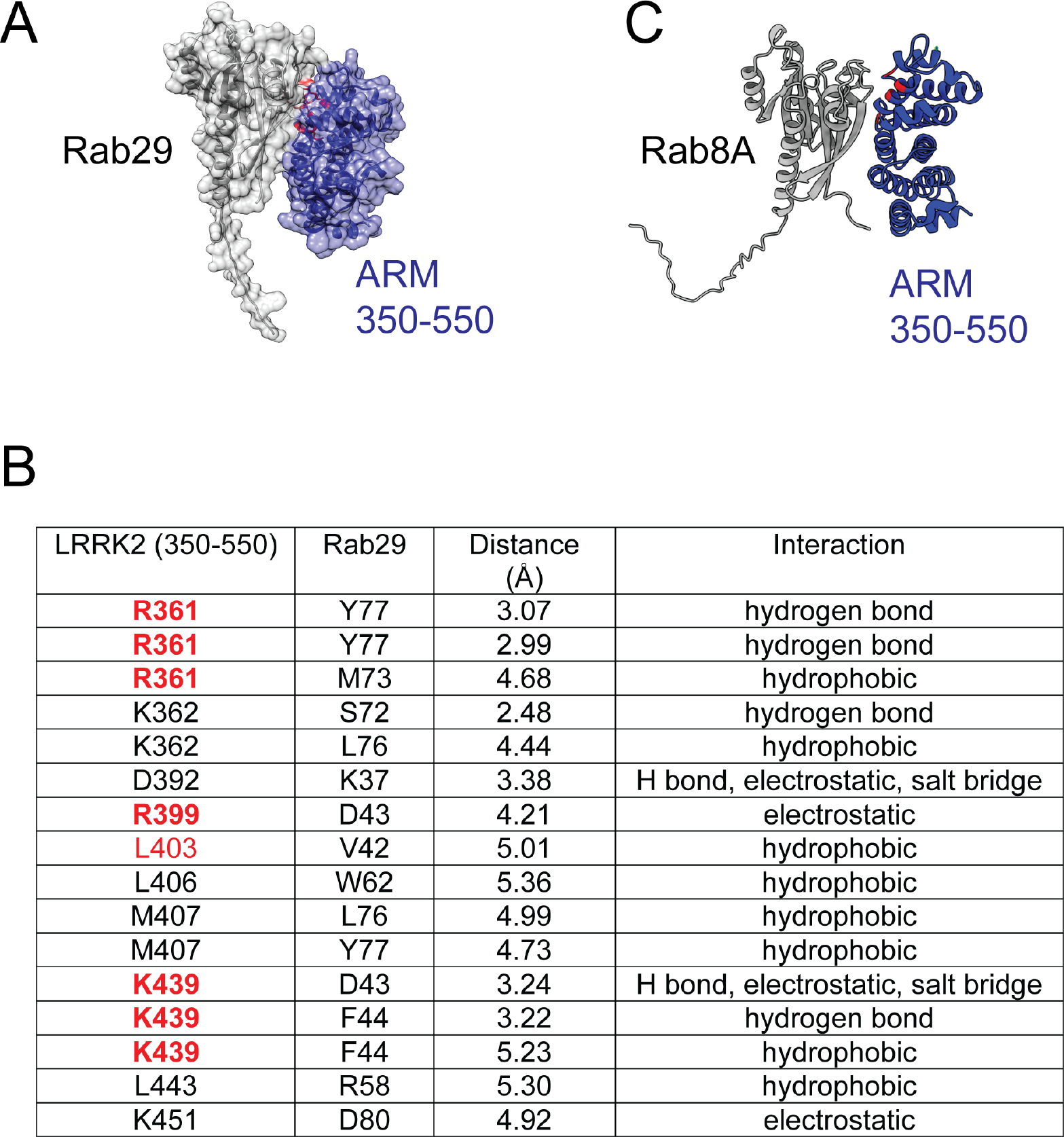
(A) Alphafold model of a complex of Rab29 (grey) bound to the 350-550 fragment of LRRK2 (purple) as in Fig. 3A. (B) Table of Site #1 residues predicted to lie within the interface of LRRK2 and Rab29. Highlighted in red are residues that when mutated, suppress interaction of LRRK2 with Rab29 and inhibit Rab29-mediated LRRK2 activation in cells. (C) Alphafold model of a complex of Rab8A (grey) bound to the 350-550 fragment of LRRK2 (navy); red residues are R361, R399, L403 and K439.

**Figure 3 – Figure Supplement 3.**
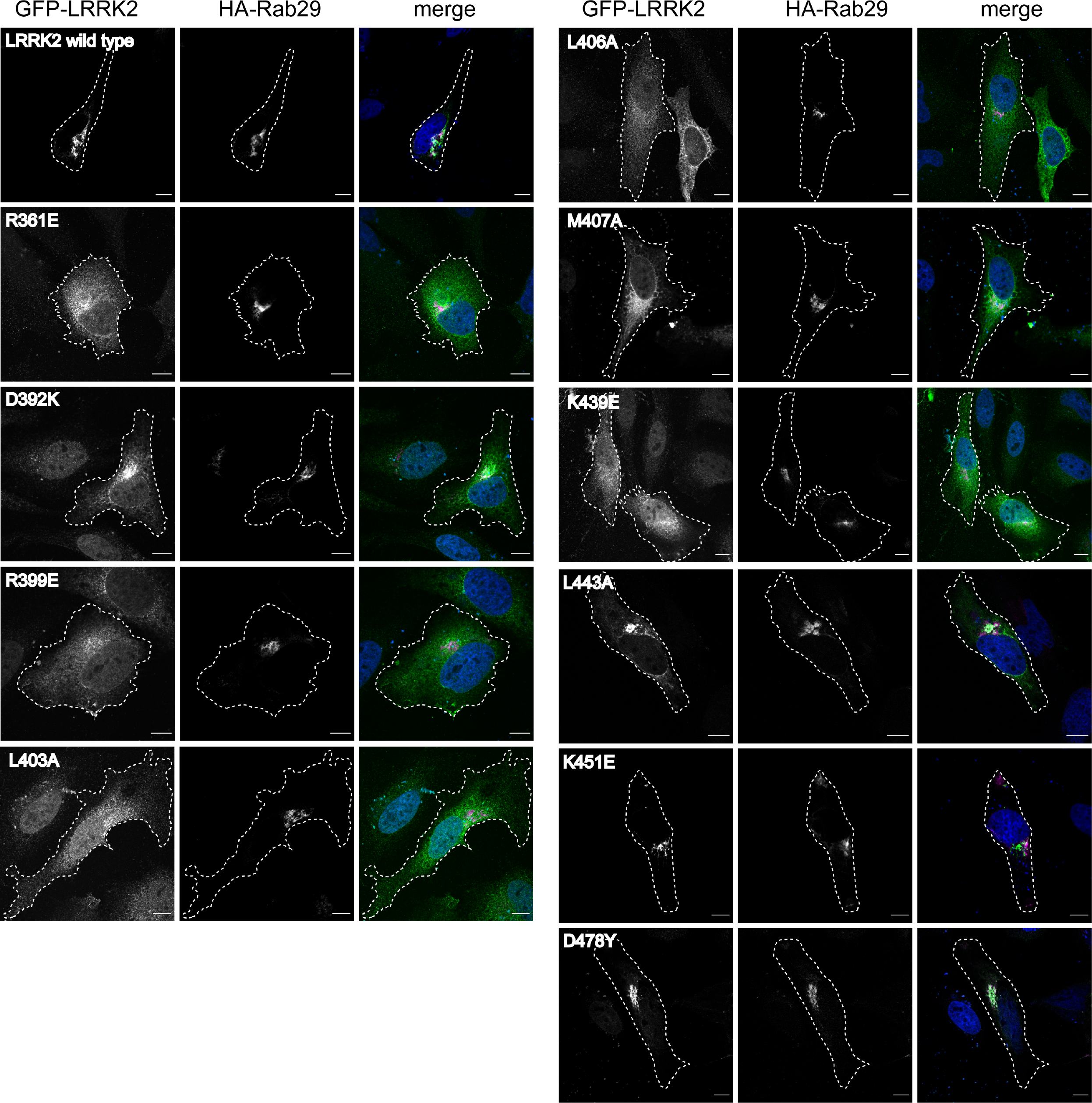
Examples of micrographs used to create Fig. 3B. Mutants (indicated) of full length GFP-LRRK2 were co-expressed with HA-Rab29 in HeLa cells. 24h post transfection, cells were fixed and localization assessed by confocal microscopy. Magnification bar, 20µm.

**Figure 3 – Figure Supplement 4.**
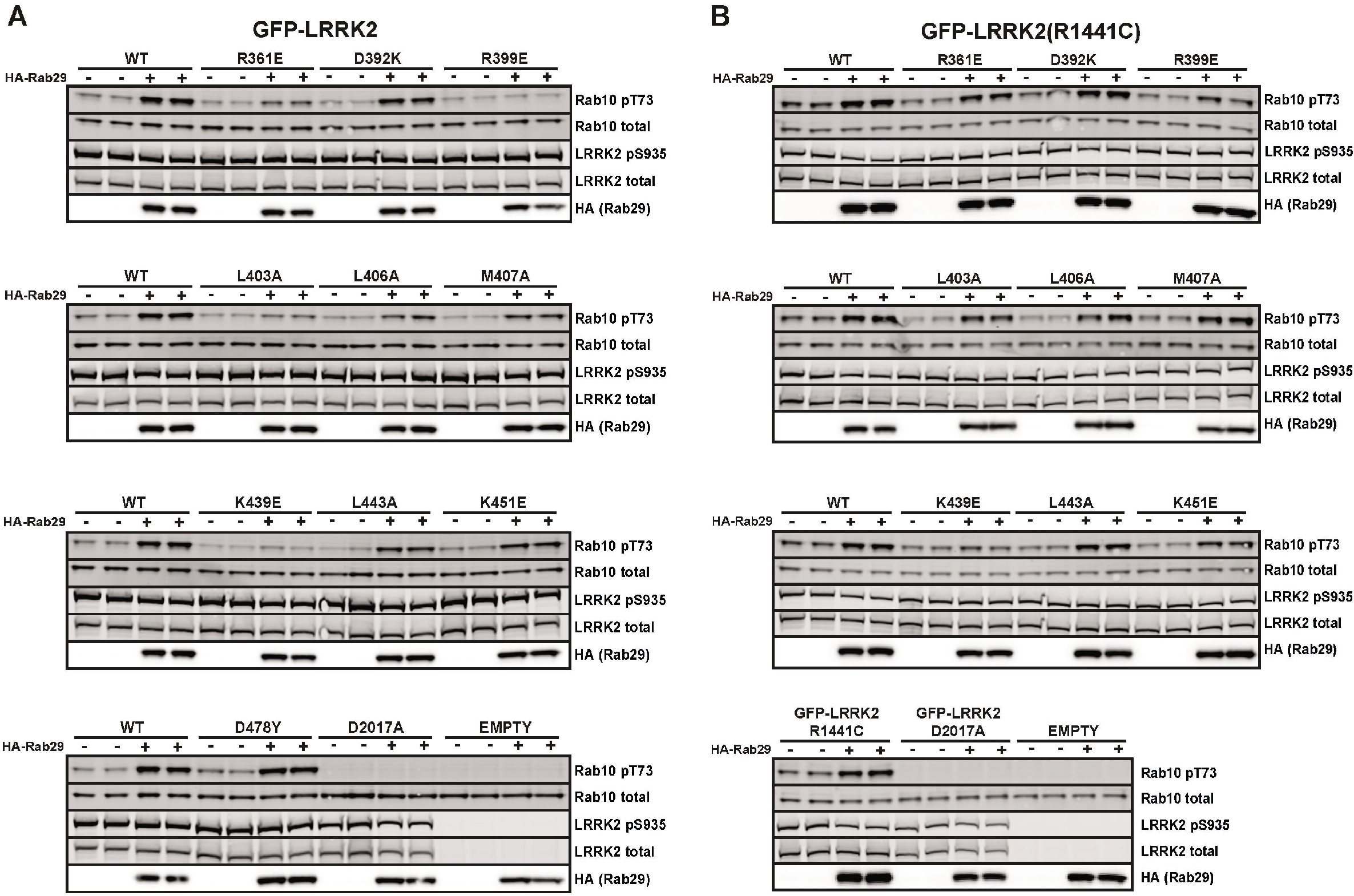
Immunoblots used to obtain Fig. 3C and Fig. 3D. Wild type and indicated mutants of full length of GFP-LRRK2 (A) or R1441G LRRK2 (B) were co-expressed with HA-Rab29 in HEK293T cells. 24h post transfection cells were fixed and samples analyzed for immunoblotting. Each membrane was probed with anti- pRab10 (rabbit), anti-Rab10 (mouse) and anti-HA (rat) antibodies. pRab10 and Rab10 signals were detected using 800 (anti-rabbit) and 680 (anti-mouse) channels in LICOR, whereas the HA (showing Rab29 expression) was developed using ECL (anti-rat).

**Figure 8 – Figure Supplement 1.**
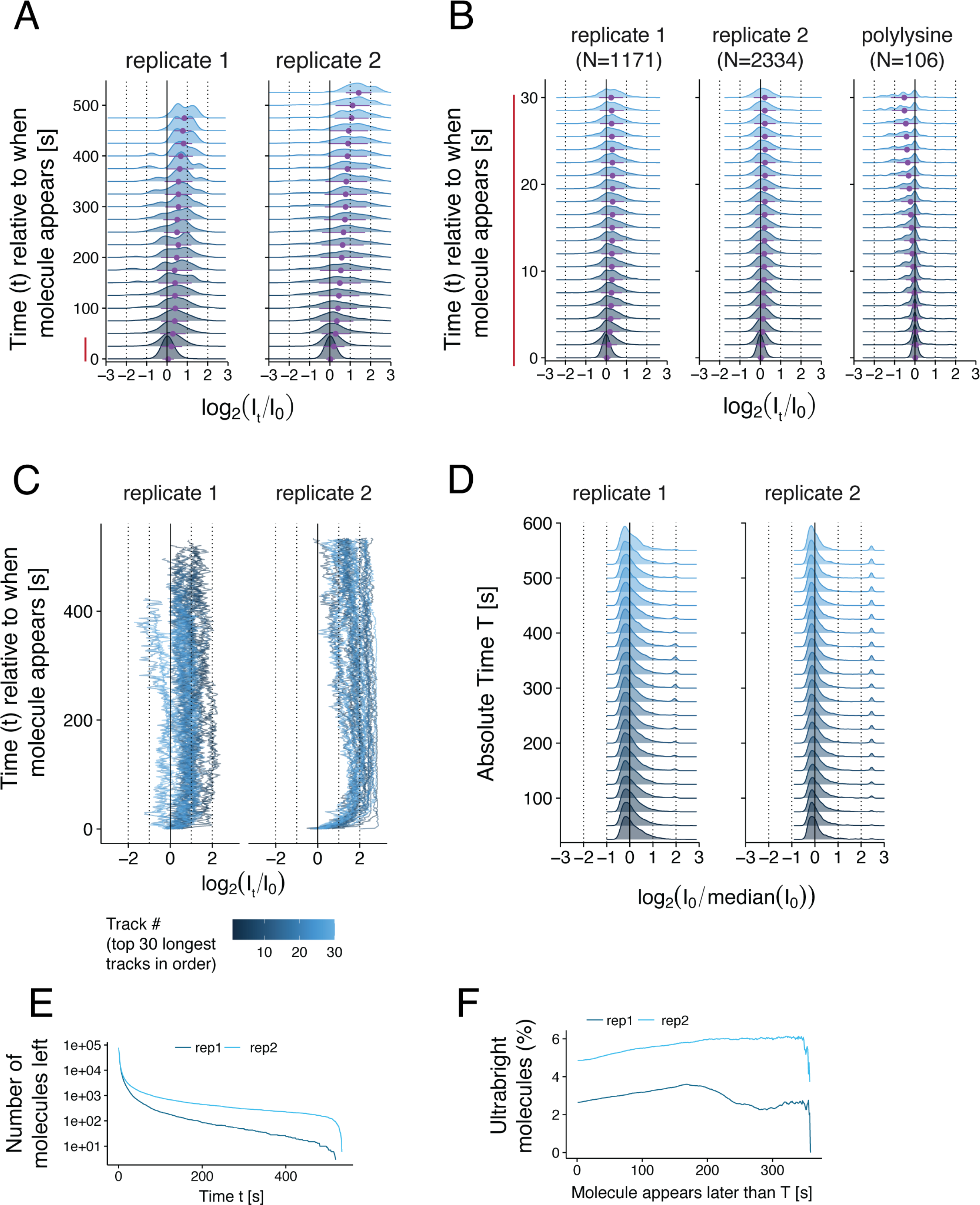
Quantitative analysis of TIRF images of LRRK2 recruitment on planar lipid bilayers. A. Ridge plot showing the distribution of the fluorescence intensity of CF633-labeled LRRK2 molecules (or complexes) as a function of the time elapsed since the molecule first appeared on the surface. Two replicates of LRRK2 are shown. For each molecule, the fluorescence intensity *I_t_* at each time point *t* was normalized with respect to its initial intensity *I_0_* during the first frame when the molecule first appeared on the surface. The intensity distributions are shown in log scale (*x* axis). Intensity distributions were computed using the average intensity of each molecule over 25s increments. Purple data points and error bars below each distribution show its mean and standard deviation. For each time point *t*, all the molecules with a fluorescence lifetime greater than *t* were used to compute the distribution. The number of such molecules at each time point is shown in panel E. B. Same ridge plot as in A showing the evolution of the fluorescence intensity with greater temporal resolution during the first 30s after a molecule appeared on the surface. Here, the intensity distributions were computed using the average intensity of each molecule over 1.5s increments. Only molecules with a fluorescence lifetime larger than 30s were included in this ridge plot (N=1171, 2334, and 106 molecules respectively for the 3 conditions shown (left to right). C. Individual fluorescence intensity over time for the 30 molecules with the longest fluorescence lifetimes as in A. D. Ridge plot showing the distribution of the initial fluorescence intensity *I_0_* of individual CF633-labeled LRRK2 molecules or complexes when they first appeared on the surface, as a function of the time elapsed since the first molecule was detected (referred to as the absolute time *T*). These initial intensities were normalized to the median initial intensity of all molecules. E. Inverse cumulative distribution of the fluorescence lifetime of individual molecules. F. Percentage of molecules that were “ultrabright” when they first appeared on the surface, as a function of the absolute time *T* at which they appeared. Ultrabright refers to molecules with an initial intensity greater than 2^1.5^ fold the median initial intensity across all molecules (log_2_[I_0_/median(I_0_)]>1.5).

**Figure 8 – Figure Supplement 2.**
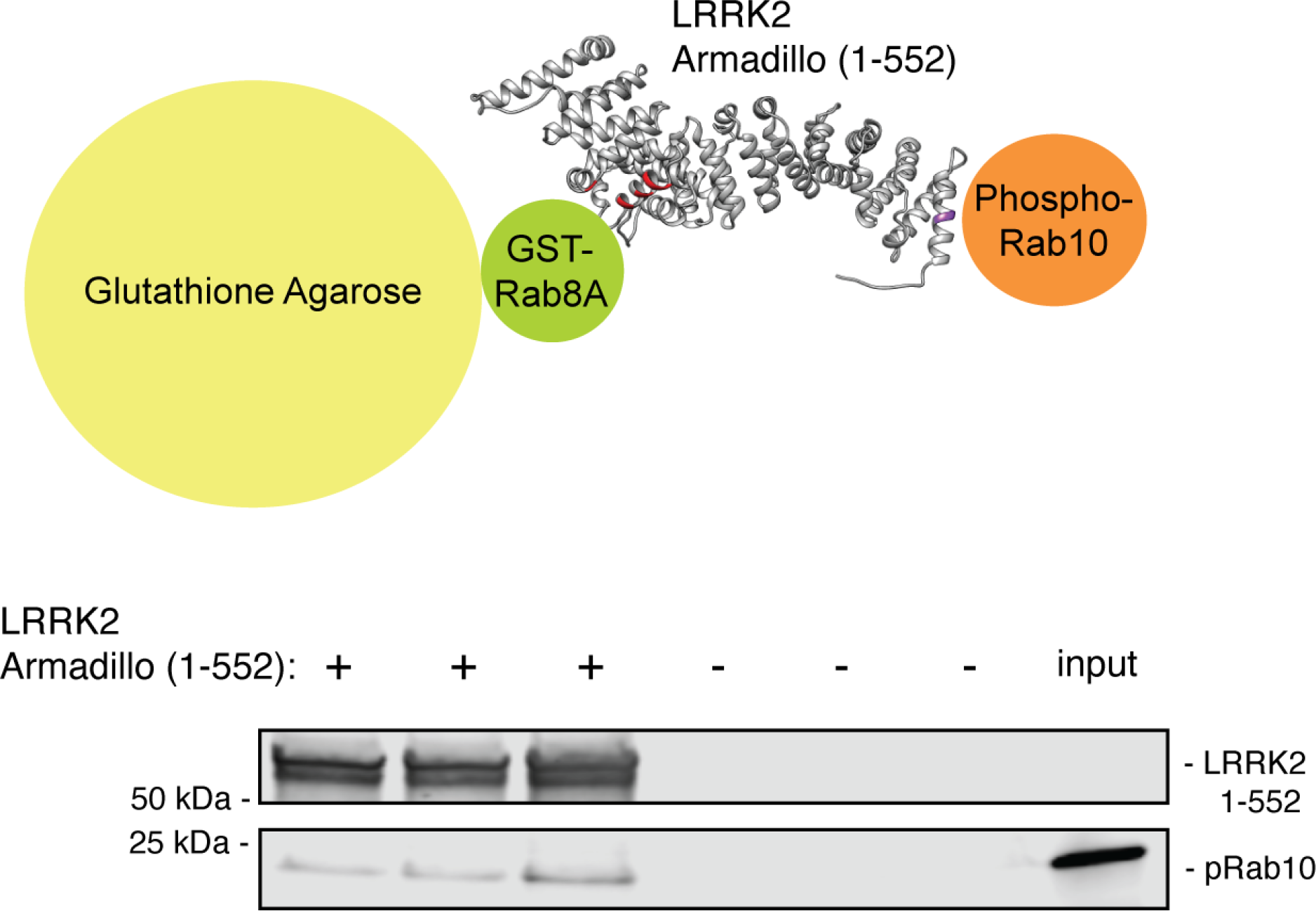
The LRRK2 Armadillo domain can bind phosphorylated Rab10 and unphosphorylated Rab8A simultaneously. GST-Rab8A was immobilized on glutathione agarose, then LRRK2 Armadillo (or buffer) was added; beads were washed and His-phosphoRab10 Q68L was then added. Bead-bound material (triplicates shown) was eluted with reduced glutathione and analyzed by immunoblotting. Input, 50% of that used in each binding reaction. PhosphoRab10 (5% of input) was detected only in ARM domain-containing samples, consistent with the Kd values.

## Supporting Videos

Video 1. TIRF microscopy of R1441G LRRK2 binding to Rab10-lipid bilayers Captured at 1 frame per second and compressed 20X

Video 2. TIRF microscopy of R1441G LRRK2 binding to lipid bilayers without Rab10 Captured at 1 frame per second and compressed 20X

Video 3. TIRF microscopy of R1441G LRRK2 binding to Rab11-lipid bilayers Captured at 0.5 frames per second and compressed 40X

Video 4. TIRF microscopy of D2017A LRRK2 binding to Rab10-lipid bilayers Captured at 1 frame per second and compressed 20X

Video 5. TIRF microscopy of K17A/K18A/R1441G LRRK2 binding to Rab10-lipid bilayers Captured at 0.5 frames per second and compressed 40X

## Source Data

Figure 3-figure supplement 4-source data 1. RAW data for gels.

Figure 3-figure supplement 4-source data 2. Annotated gels.

Figure 7-source data 1. RAW data for gels.

Figure 7-source data 2. Annotated gels.

Figure 8-figure supplement 2-source data 1. RAW data for gels.

Figure 8-figure supplement 2-source data 2. Annotated gels.

Figure 9-source data 1. RAW data for gels.

Figure 9-source data 2. Annotated gels.

